# Single-nucleus transcriptomics reveal the cytological mechanism of conjugated linoleic acids in regulating intramuscular fat deposition

**DOI:** 10.1101/2024.05.30.596608

**Authors:** Liyi Wang, Shiqi Liu, Shu Zhang, Yizhen Wang, Yanbing Zhou, Tizhong Shan

## Abstract

Conjugated linoleic acids (CLAs) can serve as a nutritional intervention to regulate quality, function and fat infiltration in skeletal muscles but the specific cytological mechanisms are still unknown. Here, we applied single-nucleus RNA-sequencing (snRNA-seq) to characterize the cytological mechanism of CLAs regulates fat infiltration in skeletal muscles based on pig models. We investigated the regulatory effects of CLAs on cell populations and molecular characteristics in pig muscles and found CLAs could promote the transformation of fast glycolytic myofibers into slow oxidative myofibers. We also observed three subpopulations including SCD^+^/DGAT2^+^, FABP5^+^/SIAH1^+^, and PDE4D^+^/PDE7B^+^ subclusters in adipocytes and CLAs could increase the percentage of SCD^+^/DGAT2^+^ adipocytes. RNA velocity analysis showed FABP5^+^/SIAH1^+^ and PDE4D^+^/PDE7B^+^ adipocytes could differentiate into SCD^+^/DGAT2^+^ adipocytes. We further verified the differentiated trajectory of mature adipocytes and identified PDE4D^+^/PDE7B^+^ adipocytes could differentiate into SCD^+^/DGAT2^+^ and FABP5^+^/SIAH1^+^ adipocytes by using high IMF content Laiwu pig models. The cell-cell communication analysis identified the interaction network between adipocytes and other subclusters such as fibro/adipogenic progenitors (FAPs). Pseudotemporal trajectory analysis and RNA velocity analysis also showed FAPs could differentiate into PDE4D^+^/PDE7B^+^ preadipocytes and we discovered the differentiated trajectory of preadipocytes into mature adipocytes. Besides, we found CLAs could promote FAPs differentiate into SCD^+^/DGAT2^+^ adipocytes via inhibiting c-Jun N-terminal kinase (JNK) signalling pathway *in vitro*. This study provides a foundation for regulating fat infiltration in skeletal muscles by using nutritional strategies and provides potential opportunities to serve pig as an animal model to study human fat infiltrated diseases.

## Introduction

Meat is one of the most important sources of animal protein for humans, and its quality is associated with human health. Recently, due to the development of economic levels and the improvement of living standards, people are seeking for ‘less but better’ meat (Sahlin & Trewern, 2022). Intramuscular fat (IMF) deposition is a key factor positively related to meat quality traits and lipo-nutritional values of meat, such as flavor, tenderness, and juiciness(Hausman, Basu, Du, Fernyhough-Culver, & Dodson, 2014; W. Yi, Huang, Wang, & Shan, 2023). Besides, fat infiltration in skeletal muscle (also known as myosteatosis) is the pathologic fat accumulation in skeletal muscle with poor metabolic and musculoskeletal health, it always accompanied by the decline of muscle quality and function(Biltz et al., 2020; Jiang, Marriott, & Maly, 2019). Myosteatosis is now considered as a common feature of ageing and is also related to some diseases(Wang, Valencak, & Shan, 2024). However, the occurrence mechanism and cell sources of fat accumulation in skeletal muscle is very complicated. Recently, with the rapid development of multi-omics including single-cell RNA sequencing (scRNA-seq)(Tabula Muris et al., 2018), single-nucleus RNA-seq (snRNA-seq)(Petrany et al., 2020), and spatial transcriptomics (ST)(Jin et al., 2021), more and more cell types have been found to contribute to lipid deposition in skeletal muscle including myogenic cells (e.g., satellite cells (SCs)(Asakura, Komaki, & Rudnicki, 2001) and myogenic factor 5 (Myf5)^+^ mesenchymal stem cells (MSCs)(Yin et al., 2013)) and non-myogenic cells (e.g., fibro/adipogenic progenitors (FAPs)(Uezumi, Fukada, Yamamoto, Takeda, & Tsuchida, 2010), fibroblasts, myeloid-derived cells(Z. Xu et al., 2020), pericytes(Farrington-Rock et al., 2004), endothelial cells (ECs)(Lang et al., 2008), PW1^+^/Pax7^-^ interstitial cells (PICs)(Mitchell et al., 2010), and side population cells (SPs)(Tamaki et al., 2002)) based on animal models. Hence, more and more researches have been focusing on exploring the regulatory mechanism of myosteatosis at the cytological levels. Besides, fat infiltration in skeletal muscle is regulated by many influential triggers, including ageing, metabolic and nonmetabolic diseases, disuse and inactivity, and muscle injury(Wang et al., 2024). Many genes and signaling pathways participate in the formation and regulation of fat infiltration in skeletal muscle (Biferali et al., 2021; Wosczyna et al., 2021). However, the specific mechanism of nutrients regulates fat infiltration in muscle is still unknown.

Nutritional regulation strategy is one of the most vital strategies to regulate lipid accumulation in skeletal muscle based on some animal models and clinic trials, including vitamins(Gilsanz, Kremer, Mo, Wren, & Kremer, 2010; L. Zhao et al., 2020), conjugated linoleic acid (CLAs)(van Vliet, Fappi, Reeds, & Mittendorfer, 2020), linseed(Wei et al., 2016), plant extract(You et al., 2023), and so on. Hence, exploring the potential nutritional strategies to regulate fat accumulation in skeletal muscle is valuable for animal production and human health. Pigs are not only an important source of animal protein in the human diet but also serve as a valuable model for human medical biology due to their similar physiological structure in terms of size, metabolic characteristics, and cardiovascular system (Groenen et al., 2012; Lunney et al., 2021). Chinese local pig species could be used as excellent animal models to study the mechanism of lipid deposition due to they have better meat quality and high IMF content. A high IMF content always results in better juiciness, tenderness, and flavor of pork. There are different myogenesis potential in neonatal skeletal muscle between Laiwu pigs and Duroc pigs(D. D. Xu et al., 2023). Our previous study has revealed the cell heterogeneity and transcriptional dynamics of lipid deposition in skeletal muscle of Laiwu pigs and we found high IMF content Laiwu pigs had lower muscle fiber diameter(Wang, Zhao, et al., 2023). Heigai pig is a model of Chinese indigenous pig breeds, which has advantages including high farrowing rate, good pork quality, and strong disease resistance(Liyi Wang et al., 2022). Specially, we have found CLAs can improve meat quality especially increase IMF content both in lean type pig breeds and fat type pig breeds(Wang, Huang, Wang, & Shan, 2021; L. Wang et al., 2022). However, although many studies have discussed the effects of CLA on IMF deposition, there are still gaps in the cytological mechanism of CLAs in regulating lipid deposition in skeletal muscle.

Here, we present a snRNA-seq dataset collected from *longissimus dorsi muscle* (LDM) of Heigai pigs after feeding CLAs supplement to allow for analysing heterogeneity of transcriptional states in muscles. We investigated the regulatory effects of CLAs on the muscle fiber type transformation and IMF deposition. We also identified the differentiation trajectories of three subclusters in adipocytes based on Heigai pig models and high IMF content Laiwu pig models. Based on the pseudotemporal trajectories analysis, we found CLAs could promote FAPs differentiate into SCD^+^/DGAT2^+^ adipocytes via regulating mitogen-activated protein kinase (MAPK) signalling pathway. This study paves a way to regulate lipid accumulation in muscles by using nutritional strategies and provides theoretical basis on using pig as an animal model to study human muscle-related diseases.

## Results

### CLAs changed cell populations and transcriptional dynamics in LDM

Our previous study discovered CLAs improved IMF content in LDM of Heigai pigs(L. Wang et al., 2022), here, we also found CLAs significantly increased TG content but significantly decreased TC content in LDM of pigs (Figure. 1A). Meanwhile, immunofluorescence staining results showed more lipid droplets in the LDM of the CLAs group (Figure. 1B). To investigate the changes of cell heterogeneity in pig muscles after CLAs treatment at the cellular level, we performed snRNA-seq from LDM tissue of Heigai pigs using the 10× Genomics Chromium platform (Figure. 1C). First, after Cell Ranger analyses, the estimated number of cells, fraction of reads in cells, mean reads per cell, median genes per cell and median UMI counts per cell in LDM were showed in Supplementary Fig. S1a. We obtained 25507 cells from 2 individual libraries, comprising 10835 cells from CON group and 11705 cells from CLA group for the downstream analysis after the quality control of snRNA-seq data (Figure. S1B). Based on the Seurat package, we used Uniform Manifold Approximation and Projection (UMAP) plots to show the different subclusters (Figure. 1D). We identified 8 different clusters in two groups of pig muscles, including myofibers (*CAPN3*), FAPs/fibroblasts (*PDGFRA*), ECs (*CD34*), adipocytes (*PPARG*), immune cells (*PTPRC*), muscle satellite cells (MuSCs) (*PAX7*), myeloid derived cells (*MRC1*) and pericytes (*RGS5*) using the expression of marker genes (Figure. 1E). Next, the percentage of these cell types showed differences in different group (Figure. 1F). Compared with the CON group, the FAPs/fibroblasts (3.39% *vs.* 8.31%), ECs (1.26% *vs.* 2.94%), adipocytes (1.74% *vs.* 2.37%), myeloid derived cells (0.93% *vs.* 2.17%) and pericytes (0.45% *vs.* 0.7%) had a higher proportion in the CLA group. However, CLA decreased the proportion of myofibers (90.46% vs. 81.1%). The top 10 most differentially expressed genes (DEGs) were showed in the heatmap and the bar plot showed the top 3 KEGG enrichment of DEGs between the 8 different cell types in LDM (Figure. 1G). Besides, violin plots displayed CLAs upregulated the expression of mature adipocyte master genes including *ADIPOQ*, *FABP4*, *PLIN1*, and *LIPE*, adipogenic marker genes including *PPARG*, *PPARA*, *CEBPA*, and *CEBPB*, and lipid metabolism-related genes including *LPL*, *ELOVL4*, *ACAA2* and *HACD2* in pig muscles (Figures. S1C-E). These results indicated the cell types in LDM of Heigai pigs had significant differences after feeding CLAs which might induce the alterations in lipid deposition.

**Figure. 1.**
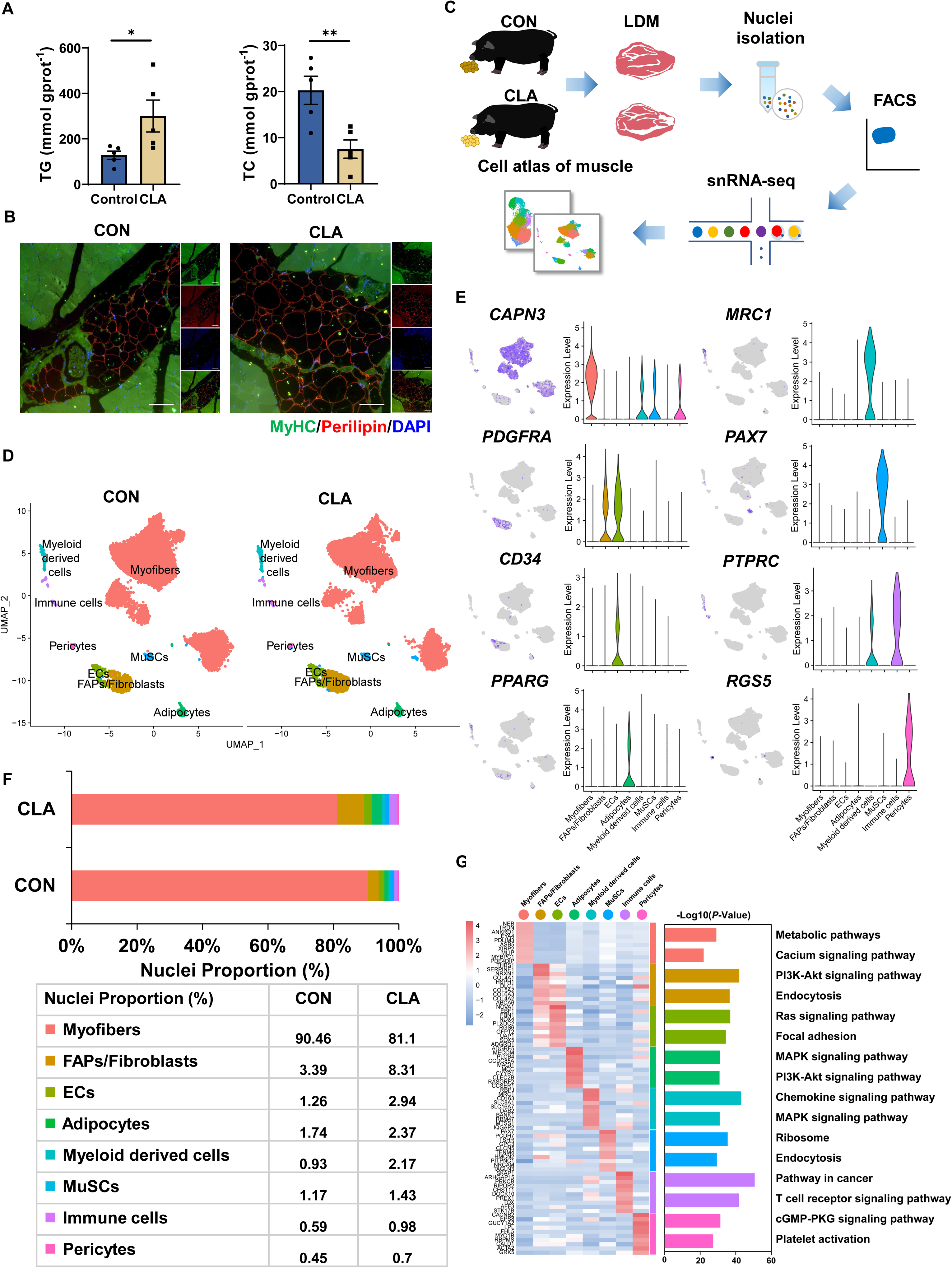
SnRNA-seq identifies distinct cell populations after CLAs treatment in pig muscles. **(A)** TG and TC content of LDM tissues in control and CLAs groups (n=5). **(B)** LDM tissues stained with the adipogenic marker perilipin (red), muscle fiber marker MyHC (green) and DAPI (blue) in different groups. Scale bars, 200 and 100 μm, respectively. **(C)** Scheme of the experimental design for snRNA-seq on different muscles. **(D)** UMAP visualization of all of the isolated single nuclei from Heigai pig muscles colored by cluster identity. **(E)** UMAP and violin plot displaying the expression of selected marker genes for each cluster in pigs. **(F)** Nuclear proportion in each cluster in pig muscles of control and CLAs groups. Each cluster is color-coded. **(G)** Left, heatmap showing the top 10 most differentially expressed genes between cell types identified. Right, KEGG enrichment for marker genes of each cell type in muscles. Each lane represents a subcluster. Error bars represent SEM. * *P* < 0.05, ** *P* < 0.01, two-tailed Student’s t-test.

### Characterization of myofibers after CLAs treatment through clustering analysis

To explore the changes of myofibers after CLA treatment, we next carried out a subcluster analysis and investigated the cell heterogeneity of myofibers. UMAP plots displayed the distribution in different subsets of myofibers (Figure. 2A). Based on our previous study(Wang, Zhao, et al., 2023) and different gene expression in myofibers, we also characterized 6 different cell types in myofibers, including I myofibers (*MYH7*), IIA myofibers (*MYH2*), IIX myofibers (*MYH1*), IIB myofibers (*MYH4*), myotendinous junctions (MTJ, *ANKRD1*), and neuromuscular junction (NMJ, *ABLIM2*) (Figure. 2B). The bar plot showed the proportion of type I myofibers (17.54% *vs.* 22.01%) and IIA myofibers (3.64% *vs.* 7.52%) had an increased tendency while the proportion of IIX myofibers (10.09% *vs.* 3.87%), IIB myofibers (63.88% *vs.* 59.35%), MTJ (6.41% *vs.* 0.28%) and NMJ (5.32% *vs.* 0.09%) had a reduced tendency in CLA group (Figure. 2C). Besides, violin plot displayed CLAs increased the expression of myofiber type marker genes (*MYH7*, *MYH2*, *MYH1*, and *MYH4*), myofiber type transformation-related genes (*PPARGC1A*, *STK11*, and *HDAC1*) and oxidation-related genes (*COX5A*, *COX5B*, and *COX8A*) but decreased glycolysis-related genes (*PFKM*, *HK2*, and *LDHC*) (Figure. 2D). Additionally, the expression of *MYHCI* and *COX5B* in LDM of Heigai pigs was also significantly increased after CLA treatment (Figure. 2E). The top 10 DEGs between the 6 cell types in myofibers were showed in heatmap (Figure. 2F). Functional enrichment analyses revealed the enrichment of metabolic pathways, oxidative phosphorylation, and thermogenesis in I myofibers and calcium, cGMP-PKG, and MAPK signalling pathway in IIB myofibers by using KEGG pathways (Figure. 2G). These data discovered the significantly heterogeneity in myofibers between two groups and CLAs could promote slow oxidative myofibers switch into fast glycolic myofibers in pig muscles.

**Figure. 2.**
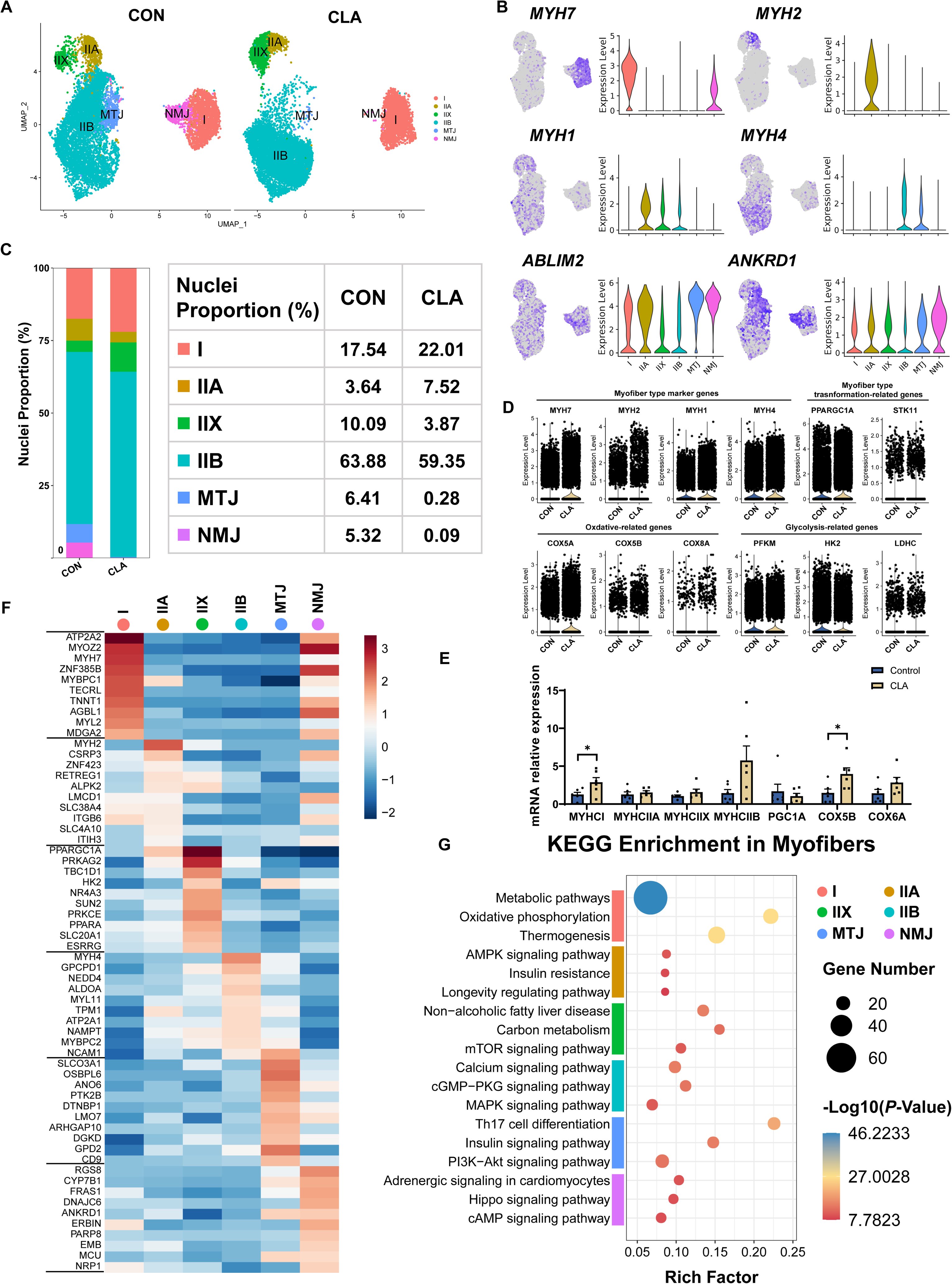
Cell and transcriptional heterogeneity in myofibers. **(A)** UMAP plot showing six subclusters of the isolated single nuclei from the control and CLAs muscles. **(B)** UMAP and violin plot displaying the expression of selected marker genes for each subcluster. **(C)** Cell proportion in each subcluster in different groups. Each cluster is colour-coded. **(D)** Violin plot showing the expression of myofiber type marker genes (*MYH7*, *MYH2*, *MYH1* and *MYH4*), myofiber type transformation-related genes (*PPARGC1A*, and *STK11*), oxidation-related genes (*COX5A*, *COX5B*, and *COX8A*), and glycolysis-related genes (*PFKM*, *HK2*, and *LDHC*) after CLAs treatment. **(E)** The mRNA expression of myofiber type related genes in LDM muscles after different treatment (n=6). **(F)** Heatmap representing the top 10 most differently expressed genes between cell subclusters identified. **(G)** KEGG enrichment for marker genes of each cell type in myofibers. I, type I myonuclei; IIa, type IIa myonuclei; IIx, type IIx myonuclei; IIb, type IIb myonuclei; MTJ, myotendinous junction nuclei; NMJ, neuromuscular junction nuclei. Error bars represent SEM. \**P* < 0.05, two-tailed Student’s t-test.

### Clustering and RNA velocity analysis revealed subpopulations and cellular dynamics of adipocytes after CLAs treatment

To investigate the regulatory mechanism of CLAs in IMF deposition, we next performed a subset analysis on adipocytes. Our previous study had discovered there are three subclusters in adipocytes nuclei of Laiwu pigs(Wang, Zhao, et al., 2023). In this study, we also characterized 3 subpopulations including SCD^+^/DGAT2^+^ adipocytes, FABP5^+^/SIAH1^+^ adipocytes, and PDE4D^+^/PDE7B^+^ adipocytes according to the most DEGs (Figures. 3A and S2A). Interestingly, we discovered CLAs increased the amounts of SCD^+^/DGAT2^+^ adipocytes (Figure. 3B) and the proportion of SCD^+^/DGAT2^+^ adipocytes (79.37% *vs.* 82.31%) but decreased the proportion of PDE4D^+^/PDE7B^+^ adipocytes (14.81% *vs.* 11.19%) (Figure. S2B). We also found CLAs increased the expression of SCD^+^/DGAT2^+^ adipocytes marker genes (*SCD* and *ARHGAP31*) and FABP5^+^/SIAH1^+^ adipocytes marker genes (*SIAH1* and *COX1*) but decreased the expression of PDE4D^+^/PDE7B^+^ adipocytes marker genes (*PDE4D*) in adipocytes (Figure. 3C). Similarly, we also discovered *SCD* and *DGAT2* expression was remarkably increased in LDM after feeding with CLAs (Figure. 3D). Meanwhile,immunofluorescence results showed more SCD1^+^ adipocytes in the LDM of the CLAs group (Figure. 3E). The top 10 most DEGs were displayed in the heatmap and the bar plot showed the top 3 KEGG enrichment of DEGs between the 3 subclusters in LDM (Figure. 3F). To explore the effects of CLAs on differentiated trajectory of adipocytes, we carried out the RNA velocity analysis of adipocytes (Figure. S2C). RNA velocity results showed the differentiated trajectory of mature adipocytes in muscles of Heigai pigs (Figure. 3G). Transcriptional dynamics of *PDE4D* and *CAPN3* were showed in Figure. S2D based on RNA velocity analysis. These results indicated in mature adipocytes, PDE4D^+^/PDE7B^+^ adipocytes and FABP5^+^/SIAH1^+^ adipocytes could differentiate into SCD^+^/DGAT2^+^ adipocytes (Figure. 3H).

**Figure. 3.**
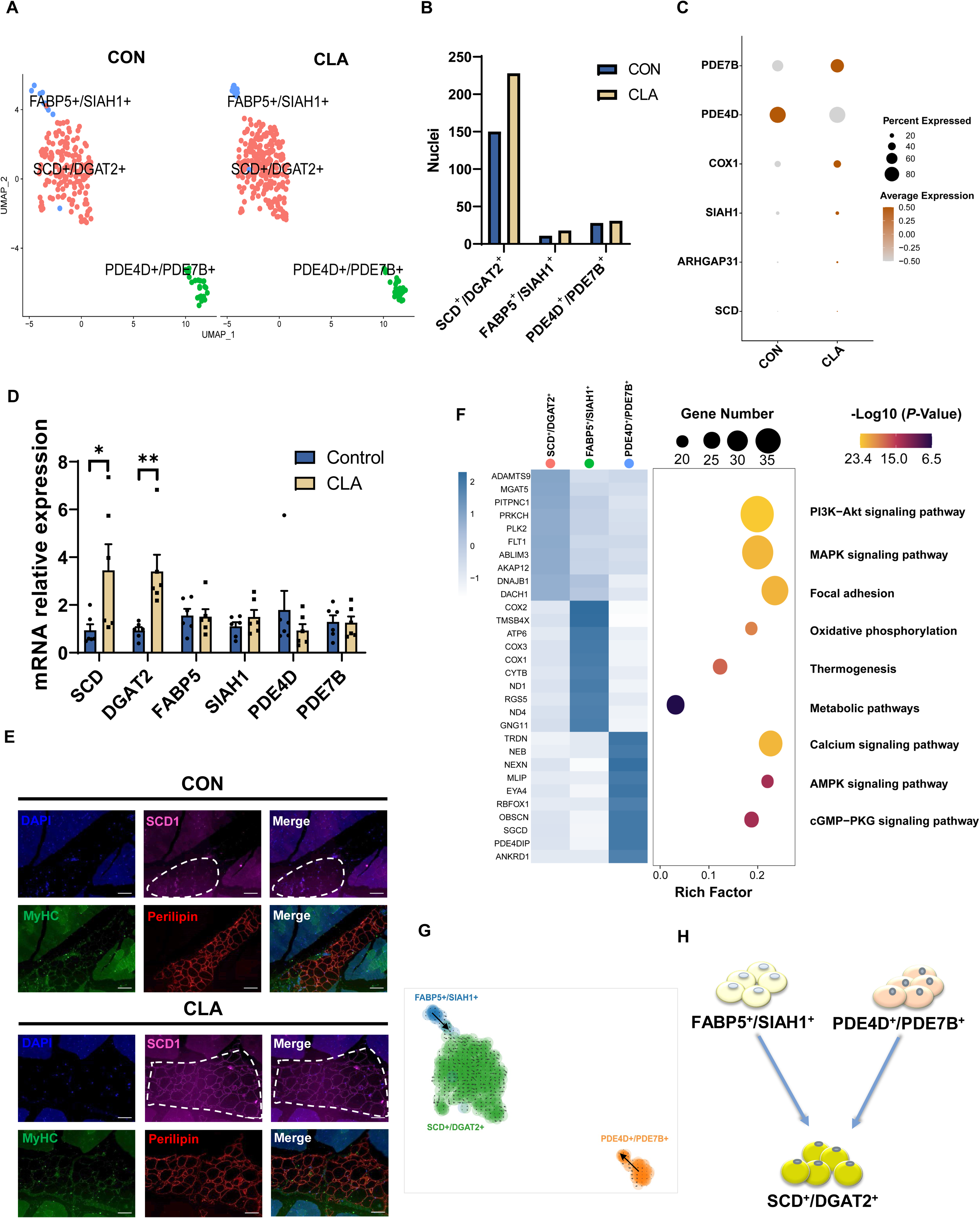
Clustering and transcriptional dynamics of adipocytes. **(A)** UMAP plot displaying the isolated single nuclei in three subclusters of adipocytes. **(B)** Bar plot displaying the cell amounts in each subcluster in different groups. **(C)** Dot plot showing the expression of three subcluster marker genes in muscle nuclei of Heigai pigs. **(D)** The mRNA expression of three subcluster marker genes in LDM muscles after different treatment (n=6). **(E)** LDM tissues stained with the adipogenic marker perilipin (red), muscle fiber marker MyHC (green), SCD1 (pink), and DAPI (blue) in different groups. Scale bars, 100 μm. **(F)** Left, heatmap showing the top 10 most differentially expressed genes between cell types identified. Right, KEGG enrichment for marker genes of each cell type in muscles. **(G)** Unsupervised pseudotime trajectory of the three subtypes of adipocytes by RNA velocity analysis. Trajectory is colored by cell subtypes. The arrow indicates the direction of cell pseudo-temporal differentiation. **(H)** Scheme of the differentiation trajectories in mature adipocytes. Error bars represent SEM. \**P* < 0.05, ** *P* < 0.01, two-tailed Student’s t-test.

### The verification of differentiated trajectories of adipocytes by using Laiwu pig models

To further verify the differentiated trajectory of adipocytes, we next carried out a pseudotemporal trajectory analysis and RNA velocity analysis of adipocytes by using our previous high IMF content pig models based on Monocle 2 and scVelo (Wang, Zhao, et al., 2023) (Figure. 4A). In adipocytes of Laiwu pigs, high IMF content Laiwu pigs (HLW) group had the higher proportion of SCD^+^ /DGAT2^+^ adipocytes but the lower proportion of PDE4D^+^/PDE7B^+^ adipocytes (Figure. 4B). Meanwhile, immunofluorescence staining results also showed the more SCD1^+^ adipocytes in HLW group (Figure. 4C). The pseudotemporal trajectory and RNA velocity analysis verified that PDE4D^+^/PDE7B^+^ subclusters could differentiate in two different directions, SCD^+^/DGAT2^+^ and FABP5^+^/SIAH1^+^ subclusters with two bifurcations (Figures. 4D-F). The pseudotemporal heatmap showed gene expression dynamics including *COX1*, *ANO4*, *PPARG*, *ADIPOQ*, *ACACA*, and *ELOVL6* at Point 2 (Figure. S3A). UMAP plots also showed transcriptional dynamics of marker genes such as *FABP4*, *ADIPOQ*, *MYBPC1*, and *EYV4* (Figures. S3B-D). Also, we discovered the expression of preadipocytes related genes including *PDGFRA*, *CD34*, *CD38*, and *WT1* was enriched in PDE4D^+^/PDE7B^+^ adipocytes and mature adipocytes related genes including *FABP4*, *ADIPOQ*, *LIPE*, *PLIN1*, *PPARG*, and *AGT* were enriched in SCD^+^/DGAT2^+^ and FABP5^+^/SIAH1^+^ adipocytes (Figure. 4G). Hence, the differentiated trajectory of mature adipocytes in muscles was displayed in Figure. 4H based on above results, which showed that PDE4D^+^/PDE7B^+^ adipocytes could differentiate into SCD^+^/DGAT2^+^ and FABP5^+^/SIAH1^+^ adipocytes, and FABP5^+^/SIAH1^+^ adipocytes also can differentiate into SCD^+^/DGAT2^+^ adipocytes. Additionally, the expression of *SCD*, *DGAT2*, *FABP5*, and *SIAH1* was upregulated but *PDE4D* and *PDE7B* expression were downregulated in HLW group (Figure. 4I). These data indicated the differentiated trajectory of mature adipocytes in muscle of pigs and the percentage of SCD^+^/DGAT2^+^ and FABP5^+^/SIAH1^+^ subclusters was higher in HLW group.

**Figure. 4.**
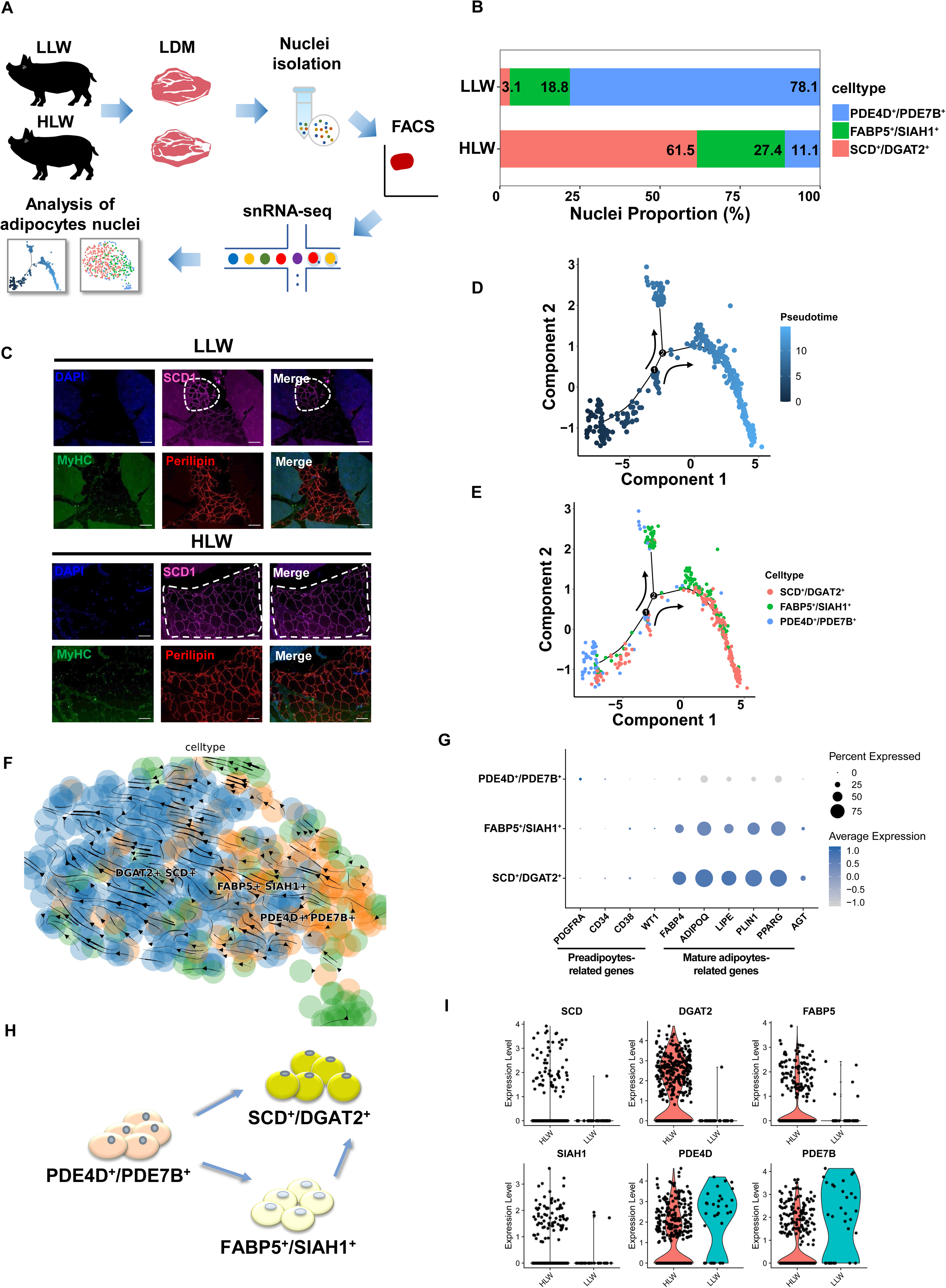
Pseudotemporal and differentiated trajectories of adipocytes in high IMF content Laiwu pig muscles. **(A)** Scheme of the experimental design for snRNA-seq on adipocytes of high IMF content Laiwu pig muscles. **(B)** Cell proportion of adipocytes subclusters in HLW and LLW groups. Each cluster is color-coded. **(C)** LDM tissues stained with the adipogenic marker perilipin (red), muscle fiber marker MyHC (green), SCD1 (pink), and DAPI (blue) in HLW and LLW groups. Scale bars, 100 μm. **(D-E)** Pseudotime ordering of all of adipocytes of subcluster DGAT2^+^/SCD^+^, FABP5^+^/SIAH1^+^, and PDE4D^+^/PDE7B^+^. Each dot represents one nucleus (color-coded by its identity), and each branch represents one cell state. Pseudotime is shown colored in a gradient from dark to light blue, and the start of pseudotime is indicated. Activation of the PDE4D^+^/PDE7B^+^ cluster can lead to DGAT2^+^/SCD^+^ and FABP5^+^/SIAH1^+^ fate. **(F)** Unsupervised pseudotime trajectory of the three subtypes of adipocytes by RNA velocity analysis. Trajectory is colored by cell subtypes. The arrow indicates the direction of cell pseudo-temporal differentiation. **(G)** Dot plot showing the expression of preadipocytes and mature adipocytes-related genes in different subclusters. **(H)** Scheme of the differentiation trajectories in mature adipocytes of Laiwu pigs. **(I)** Violin plot showing the expression of three subcluster marker genes in different groups.

### Transcriptional dynamics of glycerophospholipid metabolism in high IMF deposition pigs

Our previous studies have discovered the changes of glycerophospholipid metabolism in muscles after CLA treatment(L. Wang et al., 2022) and in high IMF content Laiwu pigs(Wang, Zhao, et al., 2023). To further investigate the transcriptional dynamics of glycerophospholipid metabolism in high IMF deposition pigs, we then compare the gene program in Heigai pigs and Laiwu pigs. After CLA treatment, the snRNA-seq dataset the expression of genes involved in the glycerophospholipid metabolism in different groups and subclusters was displayed slight differences in (Figures. S4A-C). Also, in Laiwu pigs, there are differences in gene program involved in the glycerophospholipid metabolism between two groups (Figures. S4D-F). Interestingly, we found *LCLAT1* was enriched in PDE4D^+^/PDE7B^+^ subcluster and *AGPAT3* and *AGPAT5* was enriched in SCD^+^/DGAT2^+^ subcluster (Figures. S5B and S5E). We also discovered the increase of diglycerides and phosphatidylinositols and the decrease of phosphatidic acids and phosphatidylethanolamines in high IMF deposition pigs might due to the changes of *AGPAT3*, *AGPAT4*, *AGPAT5*, *CEPT1*, and *CDIPT1* (Figures. S5C and S5F). These data revealed the significant differences in lipid composition and distribution in LDM of high IMF deposition pigs might be due to the different expression levels of glycerophospholipid metabolism-related genes.

### Cell-cell communication analysis showed the interaction between adipocytes and FAPs/fibroblasts

To further explore the association between adipocytes nuclei and other cell clusters, we applied cell-cell communication analysis by using CellPhoneDB on 8 cell types in muscles of Heigai pigs and found adipocytes mainly interacted with ECs, FAPs/fibroblasts, MuSCs, and pericytes (Figure. 5A). In addition, dot plot represented stronger communication from adipocytes to other subclusters through LRP6, FGFR1, and COL4A2 pathways in LDM of Heigai pigs (Figure. 5C). Next, we also applied cell-cell communication analysis by using CellPhoneDB on 9 cell types in muscles of Laiwu pigs and found adipocytes mainly interacted with SPs, FAPs/fibroblasts, ECs, and pericytes (Figure. 5B). Also, dot plot showed stronger communication from adipocytes to other subclusters through COL6A3, LAMC1, and THBS1 pathways in LDM of Laiwu pigs (Figure. 5D). These results suggested adipocytes has tight association with FAPs/fibroblasts.

**Figure. 5.**
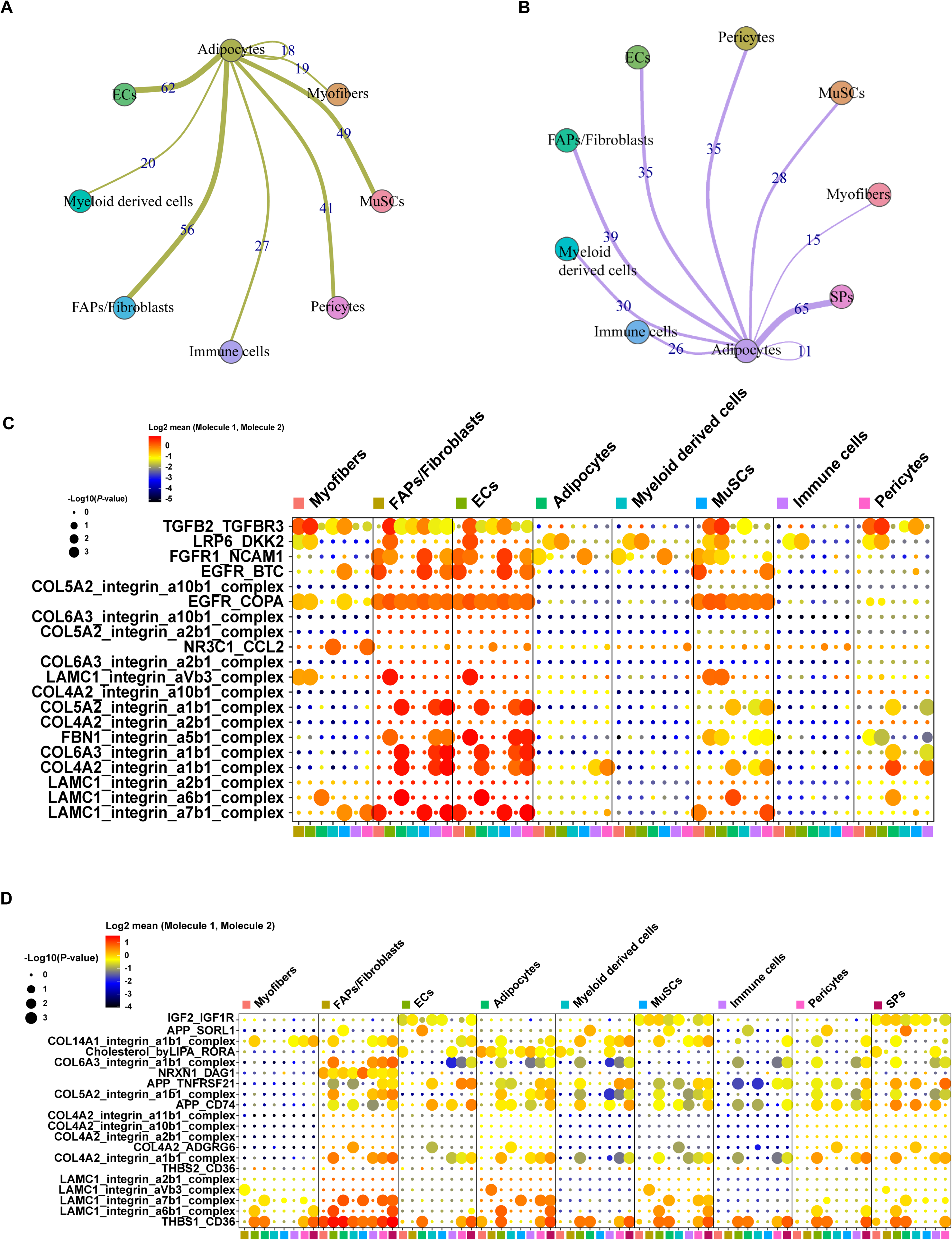
Cell-cell communication analysis of adipocytes in pig muscles. **(A)** Cell-cell communication analysis showed the network between adipocytes and other clusters in muscles of Heigai pigs. **(B)** Cell-cell communication analysis showed the network between adipocytes and other clusters in muscles of Laiwu pigs. **(C)** Dotplot representing the gene expression and significance of the receptor-ligand relationship in different cell population in muscles of Heigai pigs. **(D)** Dotplot representing the gene expression and significance of the receptor-ligand relationship in different cell population in muscles of Laiwu pigs. The larger the circle, the smaller the *P* value of the relationship in the corresponding cell population, the more significant it is.

### Characterization of FAPs after CLAs treatment through clustering and pseudotime analysis

Previous studies have demonstrated that FAPs could differentiate into mature adipocytes and are the main cell sources of IMF cells(Joe et al., 2010; Uezumi et al., 2010). To further investigate the effects of CLAs on occurrence mechanism of IMF deposition, we then performed subcluster and pseudotemporal trajectory analysis on FAPs/fibroblasts. First, UMAP plots displayed the cell distribution in different subpopulations of FAPs/fibroblasts (Figure. 6A) and we identified 3 subpopulations in FAPs/fibroblasts based on the expression of marker genes, including FAPs (*PDGFRA*), Fibroblasts (*COL1A1*), and PDE4D^+^/PDE7B^+^ subclusters (*PDE4D* and *PDE7B*) (Figure. 6B). In CLA group, we found the proportion of FAPs (70.40% *vs.* 37.06%) and PDE4D^+^/PDE7B^+^ (4.11% *vs.* 2.18%) was higher than that in CON group but the proportion of in Fibroblasts (60. 76% *vs.* 25.49%) was lower (Figure. 6C). The top 10 most DEGs between the 3 subpopulations were showed in the heatmap and the bar plot displayed the significant enrichment of the signalling pathways in muscles by using KEGG enrichment analyses and we also found calcium and cGMP-PKG signalling pathway were enriched in PDE4D^+^/PDE7B^+^ subclusters (Figure. 6D). Besides, dot plot showed CLAs upregulated the expression of preadipocytes-related genes including *CD38* and *CD34*, adipogenic master genes including *ADIPOQ*, *FABP4*, *PLIN1*, and *LIPE*, mature adipocyte marker genes including *CEBPA*, and lipid metabolism-related genes including *LPL*, *ELOVL4*, *ACAA2* and *HACD2* in FAPs/Fibroblasts (Figure. S5A). To further explore FAPs’ differentiated trajectory, we applied a pseudotemporal trajectory analysis and RNA velocity analysis of FAPs/Fibroblasts. According to the results, we found FAPs could differentiate into PDE4D^+^/PDE7B^+^ and Fibroblasts subpopulations (Figures. 6E and S5C). The pseudotemporal heatmap also displayed transcriptional dynamics including *TIMP3* and *THBS1* at Point 1 (Figure. 6F). UMAP plots also showed transcriptional dynamics of marker genes in three subpopulations such as *ITGA5*, *HSPH1*, and *ETF1* (Figures. S5B-E). To further identify the differentiated trajectory of IMF cells, we isolated primary FAPs from pigs and found the expression of *FABP4*, *ADIPOQ*, *SCD*, and *DGAT2* was significantly increased but *PDE4D* expression was significantly downregulated during adipogenic differentiation *in vitro* (Figure. 6G). Besides, the expression of *FABP5* and *SIAH1* was first significantly increased then significantly decreased during adipogenic differentiation (Figure. 6G). Hence, FAPs may first differentiate into PDE4D^+^/PDE7B^+^ preadipocytes and then differentiate into PDE4D^+^/PDE7B^+^ adipocytes (Figure. 6H). These results displayed the differentiated trajectory of preadipocytes into mature adipocytes in pig muscles and CLAs might influence this process then affect IMF deposition.

**Figure. 6.**
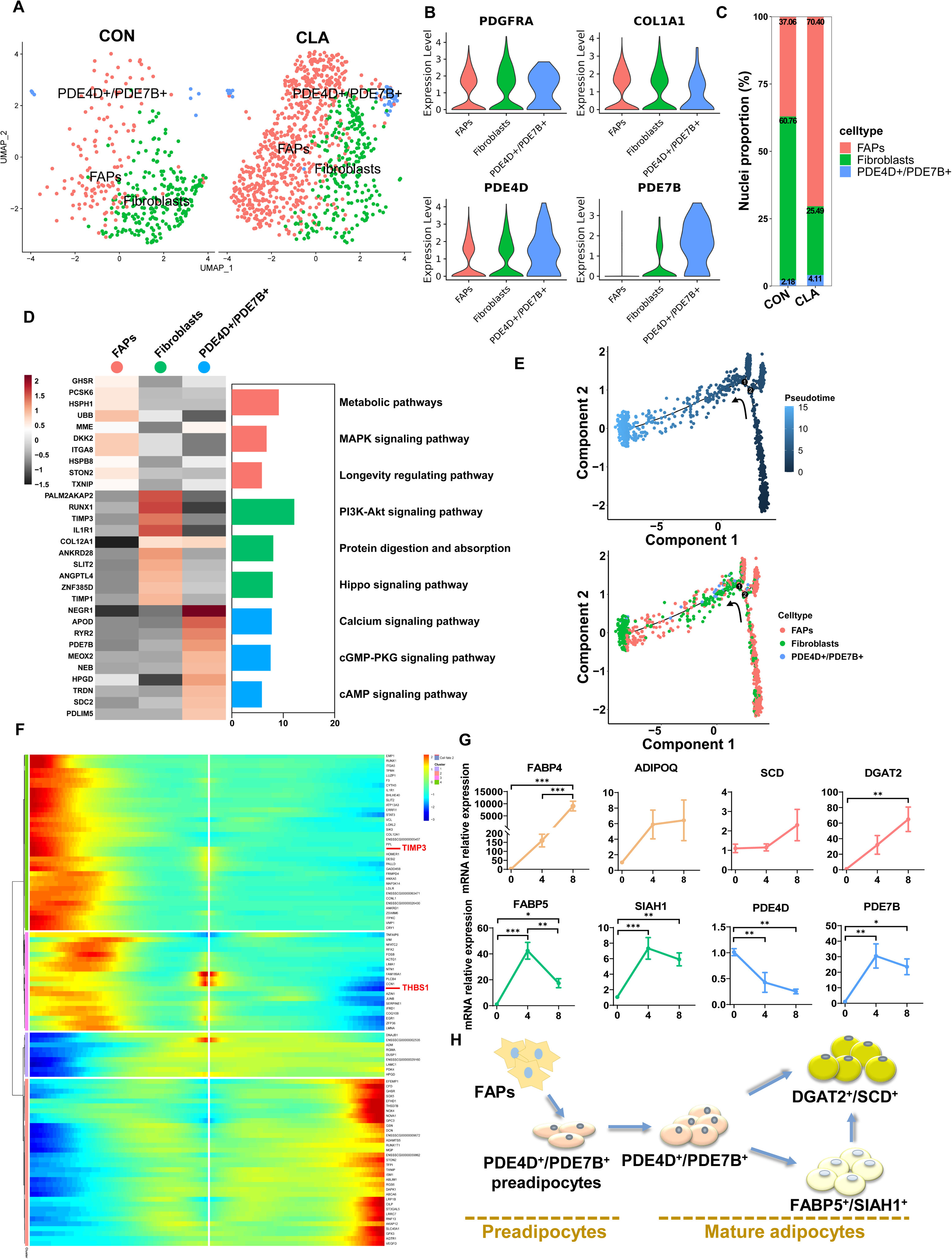
Clustering and pseudotemporal trajectories of FAPs. **(A)** UMAP plot showing three subclusters of the isolated single nuclei from control and CLAs muscle. **(B)** Violin plot displaying the expression of selected marker genes for each subcluster. **(C)** Cell proportion in each subcluster in different group. Each cluster is color-coded. **(D)** Left, heatmap showing the top 10 most differentially expressed genes between cell types identified. Right, KEGG enrichment for marker genes of each cell type in muscles. **(E)** Pseudotime ordering of all of the FAP/fibroblast of subcluster FAPs, Fibroblasts, and PDE4D^+^/PDE7B^+^. Each dot represents one nucleus (color-coded by its identity), and each branch represents one cell state. Pseudotime is shown colored in a gradient from dark to light blue, and the start of pseudotime is indicated. Activation of the FAP cluster can lead to fibroblast fate or PDE4D^+^/PDE7B^+^ fate. **(F)** Pseudotemporal heatmap showing gene expression dynamics for significant marker genes. Genes (rows) were clustered into three modules, and cells (columns) were ordered according to pseudotime in different groups. **(G)** The expression of adipogenesis and three subcluster marker genes in differentiated FAPs in different differentiation stage (n=6). **(H)** Scheme of the differentiation trajectories of preadipocytes into mature adipocytes. Error bars represent SEM. \**P* < 0.05, ** *P* < 0.01, *** *P* < 0.001, two-tailed Student’s t-test.

### CLAs promoted FAPs directed differentiation into SCD^+^/DGAT2^+^ subclusters

To further explore the cytological mechanism of CLAs regulating IMF deposition, we next investigated the regulatory effects of CLAs on the differentiation trajectory of FAPs differentiate into adipocytes. First, to explore the role of CLAs in the adipogenic differentiation of FAPs, we isolated primary FAPs from piglets and induced these adipogenic differentiation. Nile Red staining and OD490 results revealed CLAs can promote the adipogenic differentiation of FAPs after adipogenic differentiation for 8 days *in vitro* (Figures. 7A-B). Besides, the mRNA expression of *SCD*, *SIAH1* and adipogenic genes, including *FABP4*, *PPARG*, and *FASN* were significantly upregulated but *PDE4D* was significantly downregulated (Figure. 7C). The protein levels of FABP4 and SCD1 were significantly upregulated and PDE4D were significantly downregulated (Figure. 7D). In addition, the marker genes’ expression at key points of adipogenic differentiation such as *ADIPOQ*, *ELOVL6*, *ACACA*, *ARBB1*, *NEB*, and *MYBPC1* were significantly upregulated and *THBS1* and *TIMP3* were significantly downregulated (Figure. S6A). Interestingly, we next found MAPK signaling pathway were enriched in adipocytes nuclei especially SCD^+^/DGAT2^+^ subcluster (Figure. 7E). The expression of MAPK signaling pathway including ERK, c-Jun N-terminal kinase (JNK), and p38 signaling pathway related genes were changed in muscle nuclei (Figures. 7F and S6B-C). Furthermore, we observed that the protein levels of JNK phosphorylation were significantly decreased after CLA treatment during FAPs adipogenic differentiation (Figure. 7G). Hence, we next used JNK activator Anisomysin to explore the regulatory mechanism (Figure. 7H). Oil Red O staining result showed that after 24h 20 nM Anisomysin treatment, the increased adipogenic differentiation of FAPs by CLA were significantly inhibited (Figure. 7I). Also, CLA + Anisomysin group had the lower OD 490 value compared with CLA group (Figure. 7J). Nile Red staining result also showed that Anisomysin significantly inhibited the enhanced lipid droplets in FAPs after CLA treatment (Figure. 7K). MAP2K4 expression significantly upregulated and the expression of *FABP4* and *SCD* significantly decrease after Anisomysin treatment (Figure. 7L). These data indicated that CLAs may promote the directed differentiation of FAPs into SCD^+^/DGAT2^+^ subclusters via inhibiting JNK signaling pathway.

**Figure. 7.**
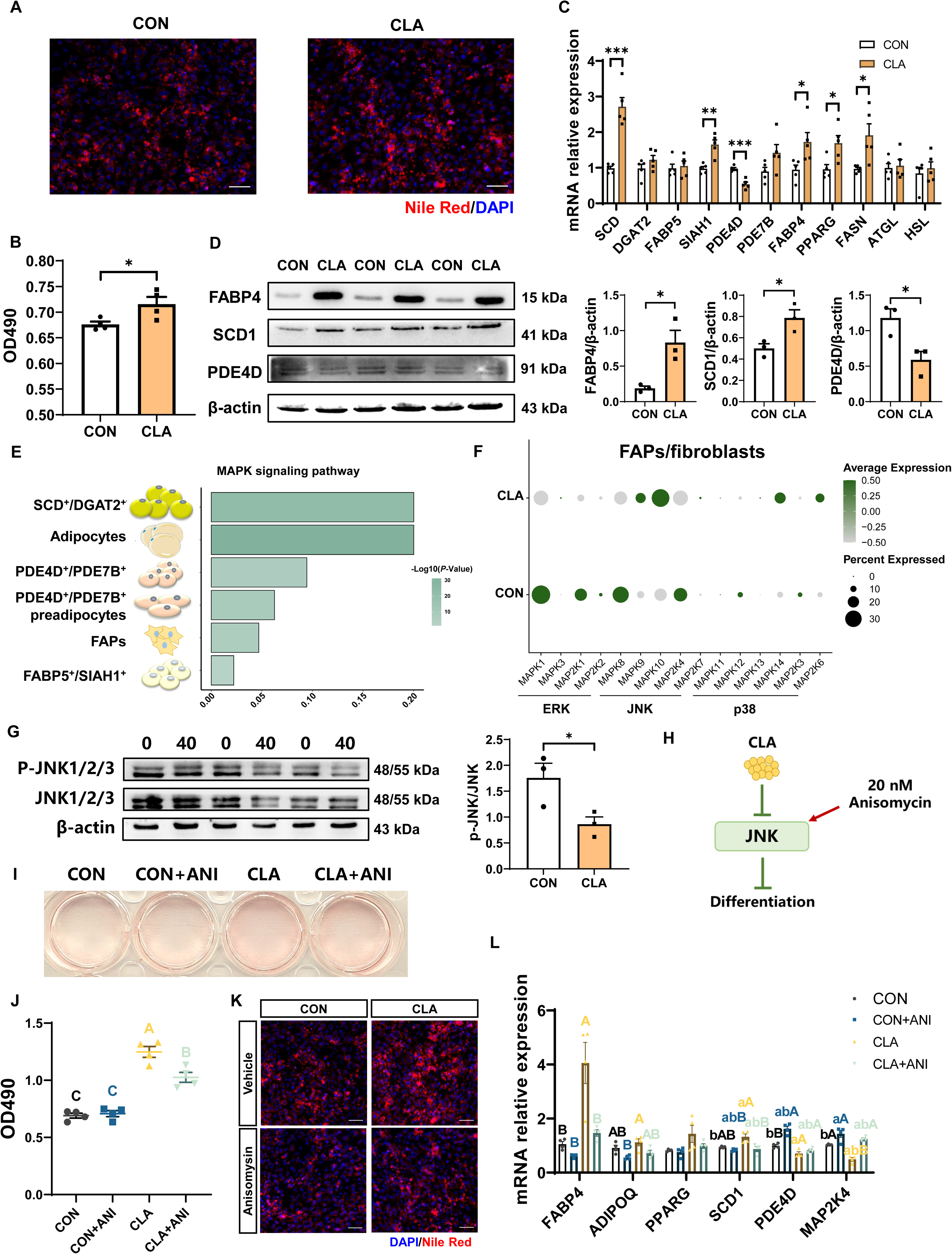
The cytological mechanism of CLAs regulates FAPs differentiation. **(A)** Dfferentiated FAPs stained with Nile Red (red) and DAPI (blue) in different groups. Scale bars, 200 and 100 μm, respectively. **(B)** OD490 levels of total lipids in differentiated FAPs after different treatment (n=4). **(C)** The mRNA expression of three subcluster marker genes and adipogenic marker genes in differentiated FAPs after different treatment (n=5). **(D)** Protein levels of FABP4, SCD1, and PDE4D were detected by western blot. **(E)** MAPK signalling pathway enrichment in different cells. **(F)** Dot plot showing the expression of MAPK signalling pathway related genes after CLA treatment in FAPs/Fibroblasts. **(G)** Protein levels of P-JNK and JNK were detected by western blot. **(H)** Scheme of CLAs regulating the differentiation trajectories of FAPs into mature adipocytes. **(I)** Dfferentiated FAPs stained with Oil Red O in different groups after treating with 20 nM Anisomysin. **(J)** OD490 levels of total lipids in differentiated FAPs after different treatment (n=4). **(K)** Dfferentiated FAPs stained with Nile Red (red) and DAPI (blue) after 20 nM Anisomysin treatment. Scale bars, 200 μm. **(L)** The mRNA expression of adipogenic related genes in differentiated FAPs after different treatment (n=4). Error bars represent SEM. \**P* < 0.05, ** *P* < 0.01, *** *P* < 0.001, two-tailed Student’s t-test and one-way ANOVA analysis. Letters represent statistical significance. Lowercase letters represent *P* < 0.05 and uppercase letters represent *P* < 0.01.

## Discussion

CLAs can serve as a nutritional intervention to regulate lipid deposition in skeletal muscle of human according to clinic trials(van Vliet et al., 2020). In animal production, CLAs could regulate meat quality in pigs and cattle, especially improve IMF content (L. Wang et al., 2022; Zhang et al., 2016). These studies pointed out that the CLAs plays a vital role in regulating fat infiltration in skeletal muscle. However, to date, the cellular mechanism of CLAs regulates lipid deposition has not been studied. Here, we utilized the 10x Genomics platform to identify the cell heterogeneity and transcriptional changes in muscles after CLAs treatment based on pig models. This study revealed the effects of CLAs on cell populations and molecular characteristics of muscles and highlighted the cytological mechanism of CLAs regulates pork quality in skeletal muscle.

CLAs is always found in ruminant animals and dairy products, it is a class of positional and geometric isomers of linoleic acids with a conjugated double bond. CLAs not only has anti-cancer, anti-hypertension, anti-adipogenic, and anti-diabetic effects, but also can improve muscle function and decrease body fat percentage. For example, adding 3.2 g/day CLAs significantly increased muscle mass in higher body fat percentage Chinese adults (Chang et al., 2020). After 0.9 g/day CLAs supplementation, body weight variation and muscle mass significantly increased and body fat percentage variation decreased in student athletes (Terasawa, Okamoto, Nakada, & Masuda, 2017). LC-MS metabolomics results discovered CLAs changed 57 metabolites which enriched in lipids/lipid-like molecules in plasma of humans (He et al., 2022). However, another study found that for sedentary older adults, CLAs had no significant influence on muscle anabolic effects (van Vliet et al., 2020). Therefore, the specific influences of CLAs on skeletal muscle is still disputed and the function on lipid deposition in human skeletal muscle needs further investigation. In animal models, many studies have demonstrated the important effects of CLAs on regulating fat accumulation in skeletal muscles. In porcine models, our foregoing studies have discovered that adding CLAs into the pig diet could significantly increase IMF contents in LDM of lean pig breeds and Heigai pigs(Wang et al., 2021; L. Wang et al., 2022). Zhang *et al* (Zhang et al., 2016)found 2% dietary CLAs significantly increased IMF deposition and reduced subcutaneous fat deposition in cattle. However, in mouse model, 0.5% mixed isomer CLAs did not lead to lipid accumulation in muscle of mice(M, S, & M, 2013). Hence, CLAs supplementation positively affect IMF deposition in muscles of pigs and ruminants but the effects of CLAs may have species-specific. In our study, we identified 8 cell types in skeletal muscles of Heigai pigs, including myofibers, FAPs/fibroblasts, ECs, adipocytes, immune cells, MuSCs, myeloid derived cells and pericytes. Recently, a variety of single cell studies have been performed on mouse/human skeletal muscles and they also identified some cell populations such as myofibers, FAPs/fibroblasts, ECs, adipocytes, immune cells, MuSCs, myeloid derived cells, tenocytes, SPs, and pericytes(Petrany et al., 2020; D. D. Xu et al., 2023; Z. Xu et al., 2020). We also found CLAs improved TG content and increased the percentage of adipocytes in LDM. Previous study demonstrated there are three subclusters in adipocytes and the formation and deposition of IMF mainly relied on DGAT2^+^/SCD^+^ adipocytes and FABP5^+^/SIAH1^+^ adipocytes(Wang, Zhao, et al., 2023). Specially, we found CLAs enhanced the percentage of SCD^+^/DGAT2^+^ subclusters. These indicated CLAs improve IMF deposition might through increasing SCD^+^/DGAT2^+^subpopulations of adipocytes. Our findings could provide a foundation for using nutritional strategies to increase pork quality especially IMF deposition. However, the specific function of CLAs on fat infiltration and deposition of skeletal muscle in people and rodents needs further study.

Skeletal muscle contains slow and fast muscle fibers and there are four major muscle fiber types in mice, including slow muscle fibers with type I muscle fibers (*Myh7*), fast muscle fibers with type IIA muscle fibers (*Myh2*), type IIX muscle fibers (*Myh1*), and type IIB muscle fibers (*Myh4*)(Dos Santos et al., 2020; Petrany et al., 2020). In pigs, illustrated there are three myofiber composition including type I, type IIA, and type IIB in skeletal muscles by ST technology(Jin et al., 2021). In this study, we identified six different subpopulations in myofibers including I myofibers, IIA myofibers, IIX myofibers, IIB myofibers, MTJ, and NMJ and IIB myofibers had the highest percentage in pig muscles. MTJ are known to exhibit structural specialization; NMJ are responsible for formation and maintenance of the synaptic apparatus and previous studies have identified these cell populations in murine skeletal muscles by using snRNA-seq(Dos Santos et al., 2020; Petrany et al., 2020). We also identified MTJ and NMJ cell populations in LDM of Laiwu pigs(Wang, Zhao, et al., 2023). These results indicate it also exist MTJ and NMJ cell populations in porcine skeletal muscles. In our previous study, we also found the percentage of type IIa myofibers had an increased tendency and type IIb myofibers had a decreased tendency in high IMF content Laiwu pigs(Wang, Zhao, et al., 2023). These results suggest that IMF content is closely related to muscle fiber type. Besides, slow muscle fibers always called slow-twitch oxidative muscle fibers like I myofibers have higher activities in mitochondrial oxidative metabolic enzymes and myoglobin while fast muscle fibers always called fast-twitch glycolytic muscle fibers like II myofibers have higher levels of glycolytic enzymes and glycogen (Schiaffino & Reggiani, 2011). Similarly, we also found I myofibers enriched in metabolic pathways, oxidative phosphorylation, and thermogenesis. In recent years, studies have focused on exploring the influences of CLAs on regulating muscle fiber type. In commercial pigs, the MyHC I mRNA abundance were improved in LDM of the CLAs group (Men, Deng, Xu, Tao, & Qi, 2013). In mice, t10, c12-CLAs, but not c9, t11-CLAs can increase oxidative skeletal muscle fiber type in gastrocnemius muscle and C2C12 myoblasts (Duan et al., 2021). CLAs have been found to prevent sarcopenia by maintaining redox balance during aging, actively regulating mitochondrial adaptation, improve muscle metabolism, and inducing hypertrophy of type IIX myofibers after endurance exercise(Barone et al., 2017; Chen, Yang, & Park, 2018). Also, we discovered CLAs enhanced the percentage of I and IIA myofibers but reduced the percentage of IIB myofibers. Previous study has found PPARγ coactivator-1α (PGC1α) serves a valuable role in skeletal muscle metabolism and is a master regulator of oxidative phosphorylation genes and could regulate muscle fiber type transformation(Handschin & Spiegelman, 2011). In our study, the PGC1α expression was also increased after CLA treatment in myofiber. These results suggested CLAs can promote glycolytic skeletal muscle fiber types switching into oxidative skeletal muscle fiber types through upregulating PGC1α expression.

Numerous studies have discovered that the cell sources of IMF cells and found several cell subsets lead to the ectopic IMF formation and deposition including SCs, Myf5^+^ MSCs, FAPs, ECs, pericytes, Fibroblasts, myeloid-derived cells, SPs, and PICs (Sciorati, Clementi, Manfredi, & Rovere-Querini, 2015; Z. Xu et al., 2020). In this study, we found CLAs increased the percentage of preadipocytes such as FAPs, ECs, myeloid-derived cells, and pericytes. Importantly, FAPs are the major source of IMF cells(Joe et al., 2010; Uezumi et al., 2010) and our previous study also verified the adipogenic capacity of FAPs in 2D and 3D culture models(Wang, Zhao, et al., 2023). Xu *et al*. found that FAPs serve as a cellular interaction hub in skeletal muscle of pigs(D. D. Xu et al., 2023). We used pseudotemporal trajectory and RNA velocity analysis combined with *in vitro* study to investigate that FAPs could first differentiate into PDE4D^+^/PDE7B^+^ adipocytes and then differentiate into DGAT2^+^/ SCD^+^ and FABP5^+^/SIAH1^+^ adipocytes. However, the regulatory mechanism of FAPs directional differentiation still needs to be further explored. *In vitro* studies demonstrated trans-10, cis-12 CLAs inhibited skeletal muscle differentiation in C2C12 cells and inhibited 3T3-L1 adipocyte adipogenesis (Hommelberg et al., 2010; Yeganeh, Taylor, Poole, Tworek, & Zahradka, 2016). However, the influences of CLAs on regulating the adipogenic differentiation of FAPs are still unclear. In this study, we found CLAs facilitated FAPs differentiating into SCD^+^/DGAT2^+^ adipocytes. Previous studies have found mice orally treated with CLAs mixture upregulated *Scd1* expression in muscle(Parra, Serra, & Palou, 2012). Besides, SCD1 expression is modulated by mTOR signaling pathway in cancer cells(J. M. Yi, Zhu, Wu, Thompson, & Jiang, 2020; S. H. Zhao et al., 2021). Moreover, MAPK signaling pathway were enriched in adipogenic differentiation of FAPs after CLAs treatment. Previous studies have discovered JNK had negative effects on regulating the adipogenic differentiation of human mesenchymal stem cells(Jang et al., 2015) and FAPs could prevent skeletal muscle regeneration after muscle injury by ST2/JNK signaling pathways(Yamakawa et al., 2023). In this study, based on JNK signing pathway activator treatment and *in vitro* experiment, we found CLAs may promote FAPs directed differentiation into SCD^+^/DGAT2^+^ adipocytes via inhibiting JNK signaling pathway. These results may provide new targets for treating human fat infiltrated diseases by nutritional strategies. However, we did not further explore the mechanistic action and the downstream transcriptional regulators need to be discussed.

In a word, we provide detailed insights into the cytological mechanism of CLAs regulates fat infiltration in skeletal muscles based on pig models via using snRNA-seq. We analysed the effects of CLAs on the cell heterogeneity and transcriptional dynamics in pig muscles and discovered CLAs could promote glycolytic muscle fiber types switching into oxidative muscle fiber types through regulating PGC1α. We also identified the differentiation trajectories of adipocytes and FAPs. Our data also demonstrated CLAs could promote FAPs differentiate into DGAT2^+^/SCD^+^ adipocytes via inhibiting JNK signalling pathway. This study provides a new way of developing nutritional strategies to combat myosteatosis and other muscle-related diseases and also offers potential opportunities to promote the utilization of pigs as animal models to study human diseases.

## Materials and methods

### Animals and samples

The Zhejiang University Animal Care and Use Committee approved all procedures and housing (ZJU20170466). 56 Heigai pigs (average body weight: 85.58 ± 10.39 kg) were divided randomly into CON group (added 1% soyabean oil) and CLA group (added 1% CLAs) for 40 days (5 days pre-feeding period and 35 days formal test period) and the nutritional levels and the feeding process as we previously reported(L. Wang et al., 2022; Wang, Zhang, Huang, Zhou, & Shan, 2023). At the end of experiment, we collected LDM from the right side of the carcass for subsequent immunofluorescence staining, biochemical assay, and snRNA-seq analyses. For Laiwu pigs, based on the determination of IMF content in Laiwu pigs, we divided them into two groups: HLW group and low IMF content Laiwu pigs (LLW) group. The two most representative samples from each group were selected for later snRNA-seq and datasets generated from muscle of high IMF content pig samples were downloaded from the Genome Sequence Archive (Genomics, Proteomics & Bioinformatics 2021) in National Genomics Data Center (Nucleic Acids Res 2022), China National Center for Bioinformation/Beijing Institute of Genomics, Chinese Academy of Sciences (GSA: CRA011059; https://ngdc.cncb.ac.cn/gsa) as we previously discussed (Wang, Zhao, et al., 2023).

### Triglycerides (TG) and total cholesterol (TC) determination

The contents of TG and TC in LDM were determined by commercial kits (TG, E1025-105; TC, E1015-50) bought from Beijing APPLYGEN Gene Technology Co., LTD.

### Immunofluorescence staining

The paraffin section was dewaxed and immersed in pre-heated sodium citrate, then placed in a microwave oven and heated for 15 min to perform antigen retrieval. Fixed the sections that cooled to room temperature in 4% paraformaldehyde for 10 min, followed by 10 min permeated with 0.5% Triton-X100and 1 h blocked with blocking buffer (5% goat serum and 2% BSA). Sections were then incubated overnight at 4 °C with Perilipin (Abcam, ab16667, 1:500), MF20 (Developmental Studies Hybridoma Bank, 1:50), and SCD1 (HuaBio, ER1916-26, 1:500) primary antibodies. Then the primary antibody was discarded and the sections were washed three times with PBS for 5 min each time. Incubated the sections with secondary antibodies for 1h and DAPI for 5 min, then washed with PBS. Sealed cell with glycerol and used fluorescent microscope to capture images.

### LDM nuclei isolation and 10X Genomics Chromium library and sequencing

LDM nuclei isolation and 10X Genomics Chromium library and sequencing were performed by LC-Bio Technology Co., Ltd. (Hangzhou, China) as previous published paper(Wang, Zhao, et al., 2023). Briefly, nuclei of LDM samples were isolated, then homogenized and incubated for 5 min on ice. The homogenate was then filtered, centrifuged and collected. The pellet was then resuspended, washed by the buffer, incubated and centrifuged. After centrifugation, the pellet was resuspended, filtered, and counted. Single-cell suspensions were loaded onto 10X Genomics Chromium for capturing 5000 single cells, followed by cDNA amplification and library construction steps were performed. Libraries were sequenced using the Illumina NovaSeq 6000 sequencing system.

### Bioinformatics analysis

SnRNA-seq results were demultiplexed and converted to FASTQ format by using Illumina bcl2fastq software and followingly processed by the Cell Ranger. Then, the Seurat packages was used to analysis the cell Ranger output. After the quality control, 22540 cells were obtained. DoubletFinder package was used to remove doublets and Harmony package was used to performed batch correction of data integration between samples. We further used Seurat, UMAP, the FindAllMarkers function to visualize the data, find clusters, and select marker genes. Monocle 2 package was used to perform trajectory analysis and model differentiation trajectories. RNA velocity analysis was independently performed in FAPs by SAMTools and the Velocyto (Bergen, Lange, Peidli, Wolf, & Theis, 2020). CellPhoneDB package was used to cell communication analysis and make further speculations about potential cellular interaction mechanisms.

### Primary FAPs isolation, magnetic cell sorting and cell culture

Primary FAPs isolation were performed as previously described (Wang, Zhao, et al., 2023). Briefly, a piece of muscle from a 3-day-old piglet was minced, and added 5 times the volume of 0.2% collagenase type I, then digested at 37 °C for 1 hour., After filtering and centrifuging, adding red blood cell lysate to split for 5 min at 4 °C followed by incubated with a Dead Cell Removal Kit at room temperature for 15 min. Then adding CD140a antibody, incubating and centrifugating. Next, added 20 μL antibiotin microbeads, incubation and centrifugation. After passing through the magnetic column, the cells on the adsorption column were PDGFRα+ cells. For FAPs adipogenic differentiation, when the cells confluence reached 90%, the10% FBS growth medium was replaced with induction medium after 4 days, the medium was changed to differentiation mediumand cultured for another 4 days until the adipocytes were mature.

### Nile Red staining

Rinse cultured FAPs with 1 x PBS 3 times, discard PBS, fix FAPs with 4% formaldehyde for 15 min, repeat the rinsing step, and then add Nile Red solution (1:500 for lipid droplet staining) and DAPI (1:500 for nuclei staining) for 5 min. Sealed cell with glycerol and used fluorescent microscope to capture images.

### Total RNA extraction and quantitative real-time PCR (qPCR)

Total RNA extraction and qPCR were conducted as described before (Shan et al., 2016). Briefly, total RNA of FAPs were extracted by using TRIzol and the Spectrophotometer and a ReverAid First Strand cDNA Synthesis Kit were used to measure the purity and concentration of total RNA and reversed RNA samples. qPCR was performed by using Applied Biosystems StepOnePlus Real-Time PCR System with Hieff qPCR SYBR® Green Master Mix and gene-specific primers (Supplementary Table 1). Relative changes in gene expression were analysed using the 2^-ΔΔCT^ method and normalized using 18S ribosomal RNA as an internal control.

### Protein extraction and western blotting

Protein extraction and western blotting were carried out as mentioned previously (Shan et al., 2016). In brief, total proteins were isolated from cells or tissues with RIPA buffer. After measuring the concentrations, proteins were separated using SDS-PAGE and subsequently transferred to a polyvinylidene fluoride membrane (PVDF, Millipore Corporation). Then blocked PVDF membrane with blocking buffer (5% fat-free milk) for 1 h and incubated with primary antibodies overnight at 4 °C. The peroxisome proliferator-activated receptorγ (PPARγ) (C26H12, 1:1000) were purchased from Cell Signaling Technology (CST). The β-actin (M1210-2, 1:10000), FABP4 (E71703-98, 1:2000), SCD1 (ER1916-26, 1:1000), PDE4D (ER1916-26, 1:500), p-JNK1/2/3(T183+T183+T221) (ET1609-42, 1:2000), and JNK1/2/3 (ET1601-28, 1:2000) antibody were from HuaBio. Dilution of the secondary antibody was 500-fold. The ChemiScope3500 Mini-System was used for protein detected.

### Statistical analysis

GraphPad (Prism 8.3.0) was used for data analyses and R software (version 4.3.2) was used for data visualization. Data comparisons were made by unpaired two-tailed Student’s t tests and one-way ANOVA analysis. Differences were considered significant at *P* < 0.05.

## Supporting information

response to the comments

## Author contribution

**Liyi Wang:** conceptualization, data curation, formal analysis, investigation, methodology, software, validation, visualization, writing-original draft. **Shiqi Liu:** investigation, methodology, writing-review & editing. **Shu Zhang:** investigation. **Yizhen Wang:** resources, supervision. **Yanbing Zhou:** conceptualization, investigation, methodology, supervision. **Tizhong Shan:** conceptualization, funding acquisition, project administration, resources, supervision, writing-review & editing.

## Acknowledgement

We thank members of the Shan Laboratory for comments and this work was partially supported by the National Natural Science Foundation of China (32272887), the Natural Science Foundation of Zhejiang Province (LZ22C170003), and the “Hundred Talents Program” funding from Zhejiang University to TZS.

## Data availability

The raw snRNA-seq data reported in this paper have been deposited in the Genome Sequence Archive (Genomics, Proteomics & Bioinformatics 2021) in National Genomics Data Center (Nucleic Acids Res 2022), China National Center for Bioinformation / Beijing Institute of Genomics, Chinese Academy of Sciences (GSA: CRA022605) that are publicly accessible at https://ngdc.cncb.ac.cn/gsa.

## Graphical abstract

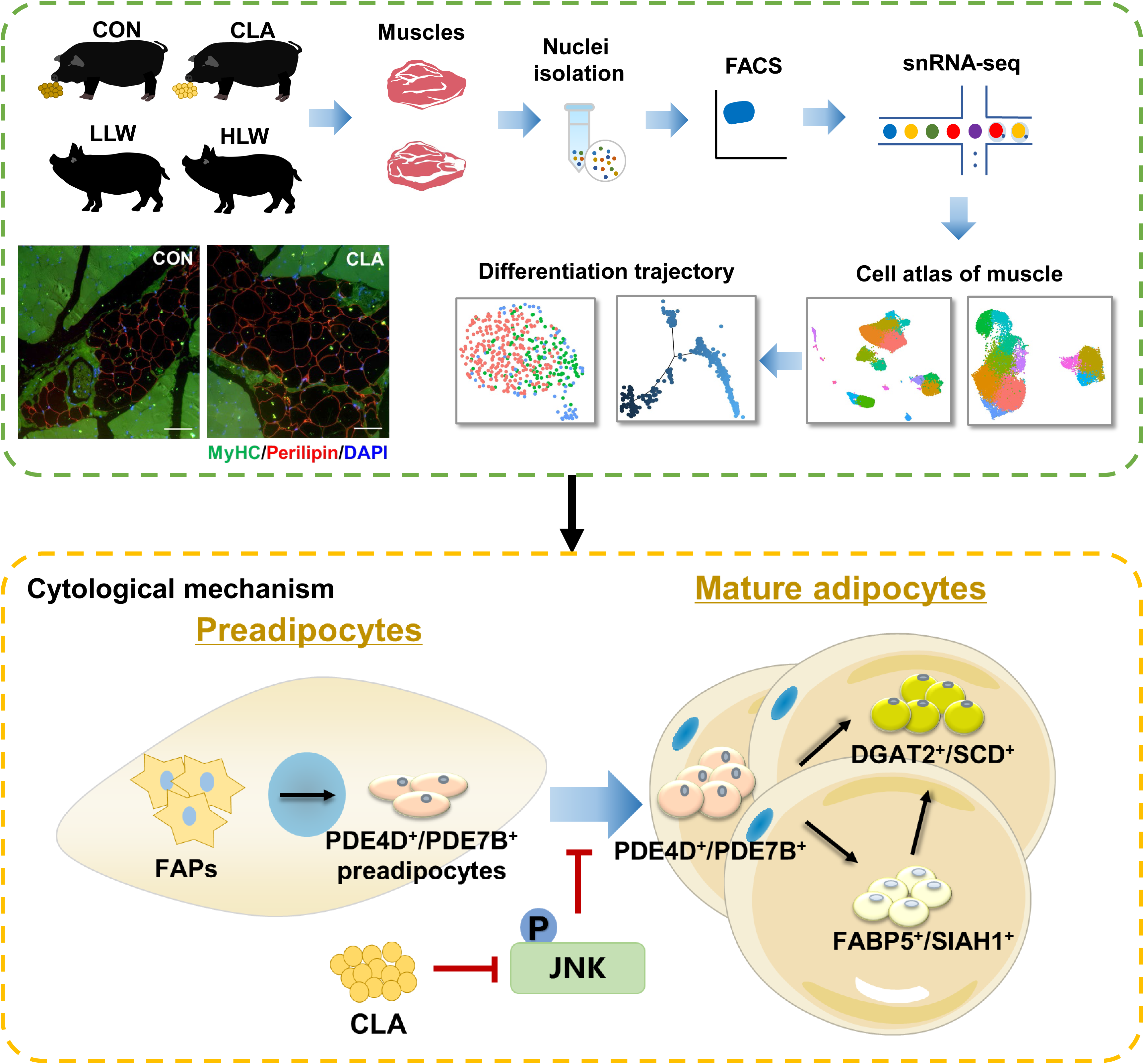

---

**Figure S1.**
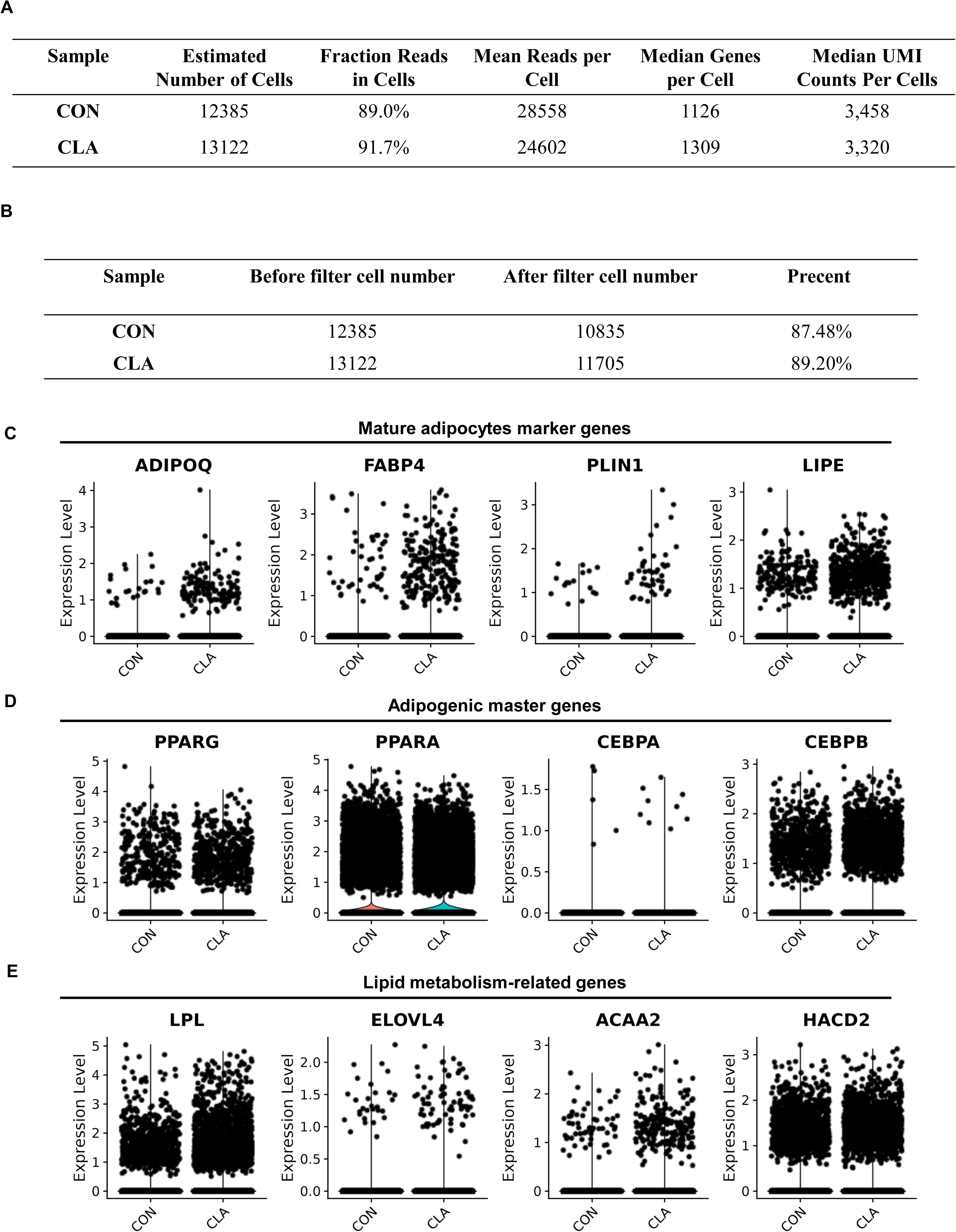

**Figure S2.**
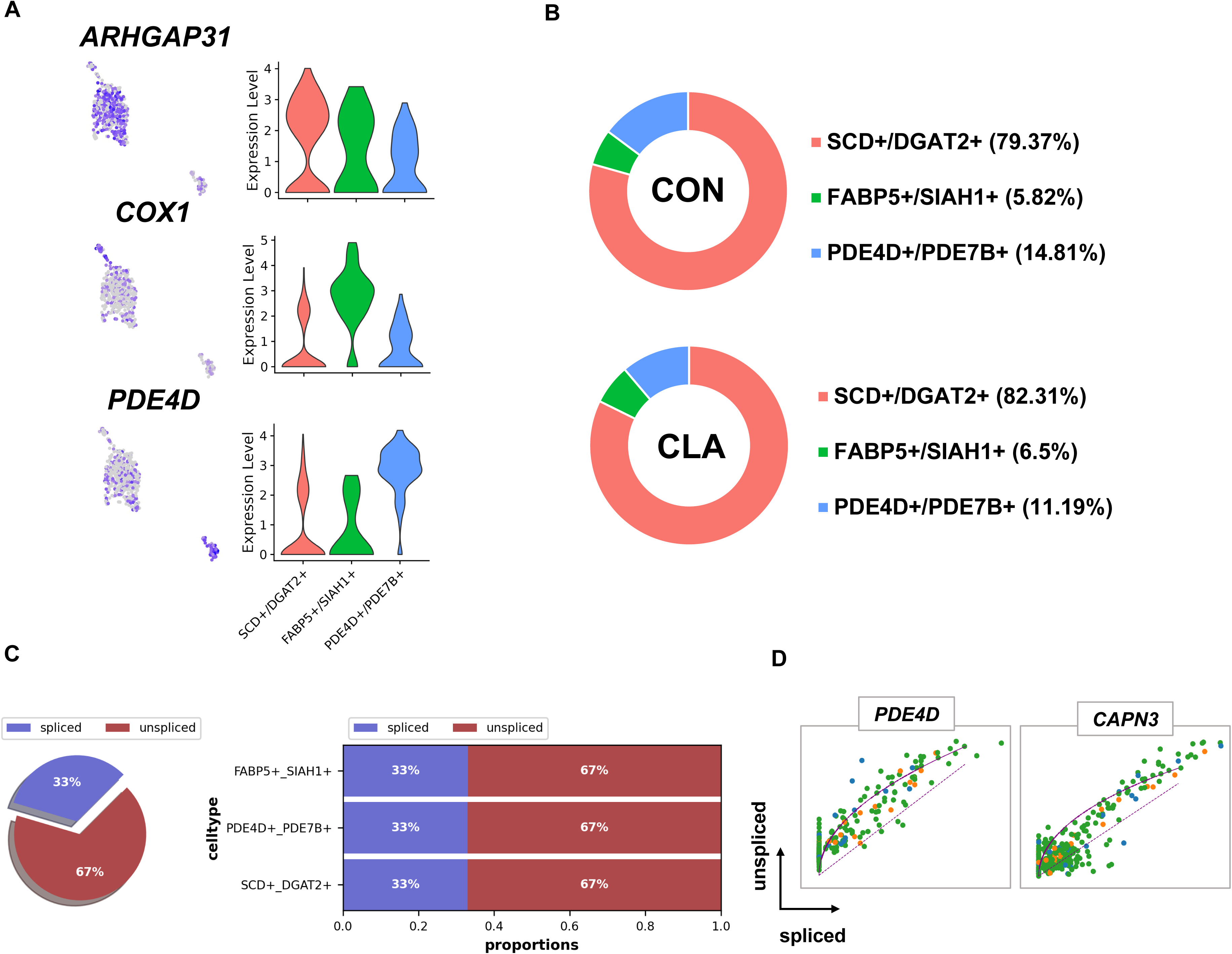

**Figure S3.**
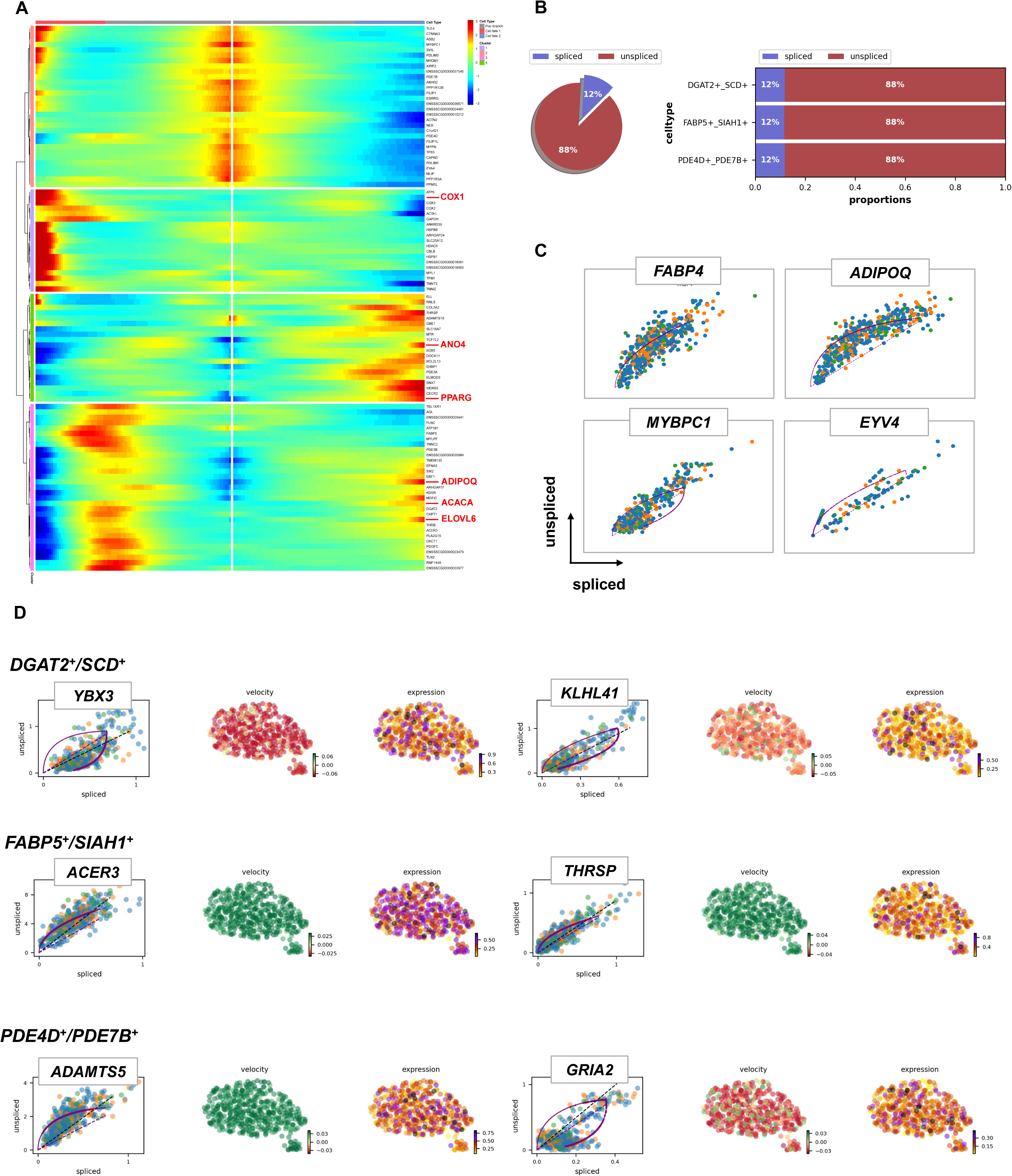

**Figure S4.**
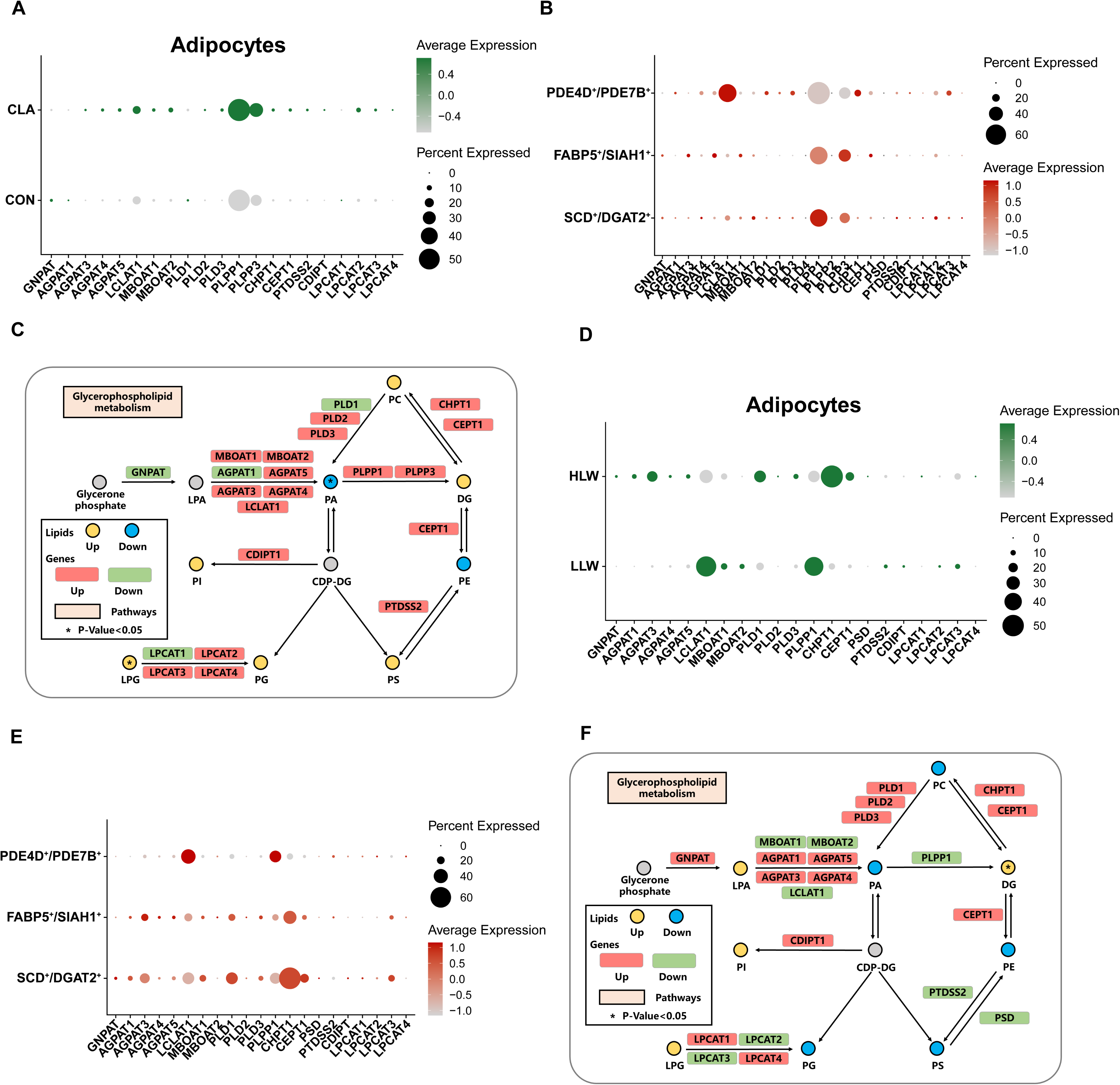

**Figure S5.**
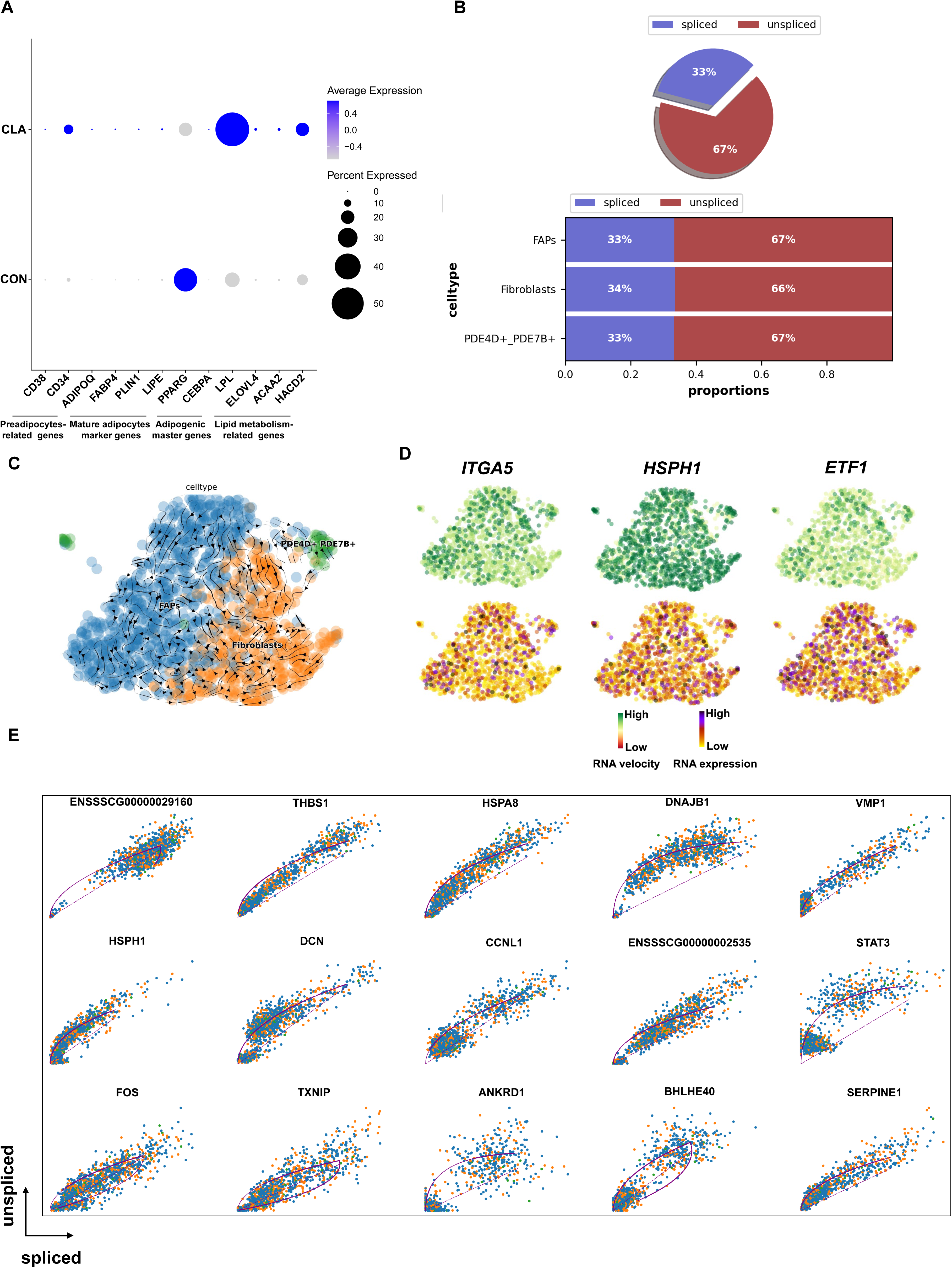

**Figure S6.**
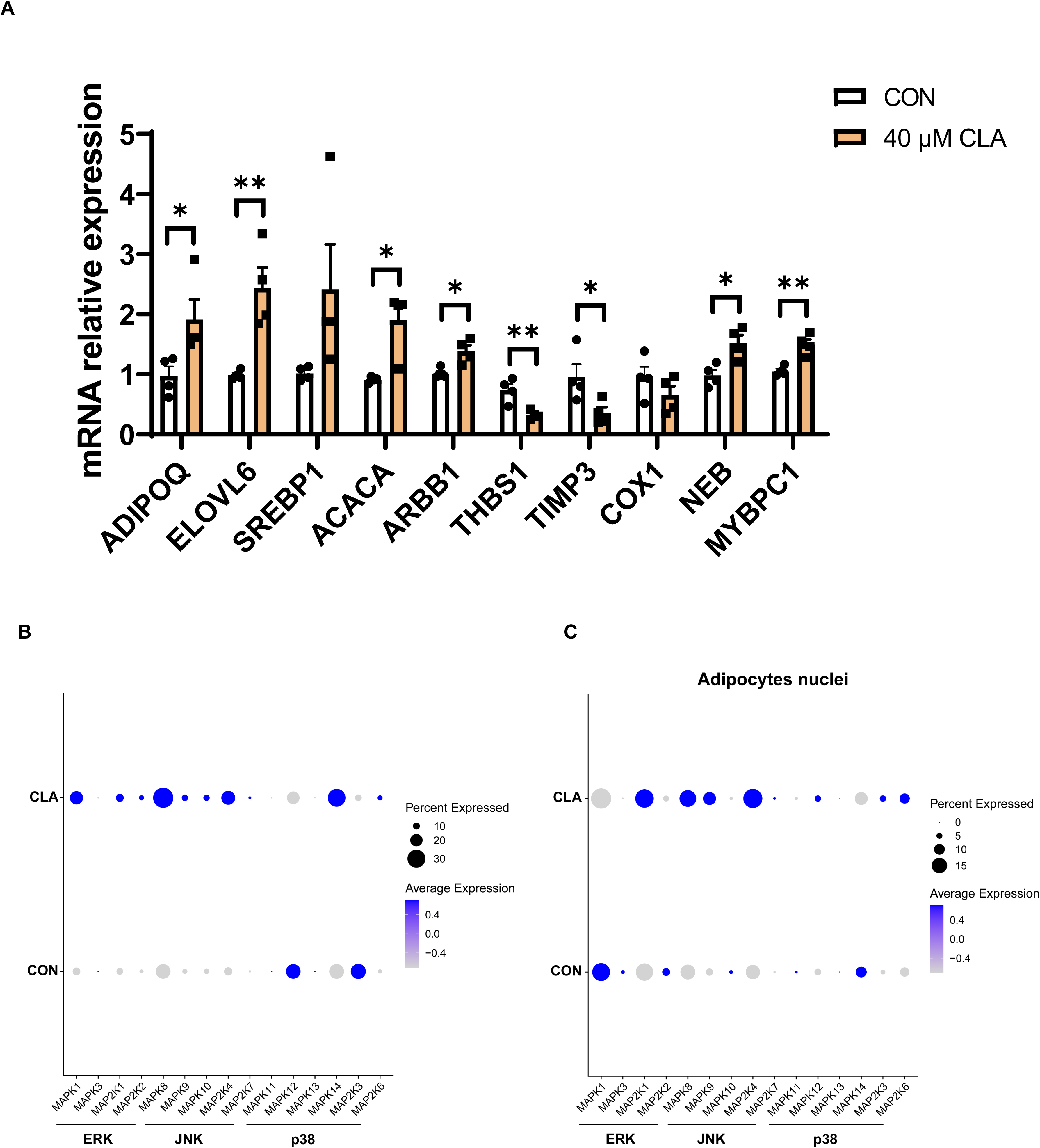

## References

Asakura, A., Komaki, M., & Rudnicki, M. A. (2001) Muscle satellite cells are multipotential stem cells that exhibit myogenic, osteogenic, and adipogenic differentiation Differentiation 68: 245-253. doi:10.1046/j.1432-0436.2001.680412.x

Barone, R., Sangiorgi, C., Marino Gammazza, A., D’Amico, D., Salerno, M., Cappello, F., . . . Macaluso, F. (2017) Effects of Conjugated Linoleic Acid Associated With Endurance Exercise on Muscle Fibres and Peroxisome Proliferator-Activated Receptor gamma Coactivator 1 alpha Isoforms J Cell Physiol 232: 1086-1094. 10.1002/jcp.25511

Bergen, V., Lange, M., Peidli, S., Wolf, F. A., & Theis, F. J. (2020) Generalizing RNA velocity to transient cell states through dynamical modeling Nat Biotechnol 38: 1408-1414. 10.1038/s41587-020-0591-3

Biferali, B., Bianconi, V., Perez, D. F., Kronawitter, S. P., Marullo, F., Maggio, R., . . . Mozzetta, C. (2021) Prdm16-mediated H3K9 methylation controls fibro-adipogenic progenitors identity during skeletal muscle repair Sci Adv 7: eabd9371. doi:10.1126/sciadv.abd9371

Biltz, N. K., Collins, K. H., Shen, K. C., Schwartz, K., Harris, C. A., & Meyer, G. A. (2020) Infiltration of intramuscular adipose tissue impairs skeletal muscle contraction J Physiol 598: 2669-2683. 10.1113/JP279595

Chang, H., Gan, W., Liao, X., Wei, J. X., Lu, M. N., Chen, H. T., . . . Liu, X. (2020) Conjugated linoleic acid supplements preserve muscle in high-body-fat adults: A double-blind, randomized, placebo trial Nutrition Metabolism and Cardiovascular Diseases 30: 1777-1784.10.1016/j.numecd.2020.05.029

Chen, P. B., Yang, J. S., & Park, Y. (2018) Adaptations of Skeletal Muscle Mitochondria to Obesity, Exercise, and Polyunsaturated Fatty Acids Lipids 53: 271–278. 10.1002/lipd.12037

Dos Santos, M., Backer, S., Saintpierre, B., Izac, B., Andrieu, M., Letourneur, F., . . . Maire, P. (2020) Single-nucleus RNA-seq and FISH identify coordinated transcriptional activity in mammalian myofibers Nat Commun 11: 5102. doi:10.1038/s41467-020-18789-8

Duan, C., Yin, C., Ma, Z., Li, F., Zhang, F., Yang, Q., . . . Wang, S. (2021) trans 10, cis 12, but Not cis 9, trans 11 Conjugated Linoleic Acid Isomer Enhances Exercise Endurance by Increasing Oxidative Skeletal Muscle Fiber Type via Toll-like Receptor 4 Signaling in Mice J Agric Food Chem 69: 15636-15648. doi:10.1021/acs.jafc.1c06280

Farrington-Rock, C., Crofts, N. J., Doherty, M. J., Ashton, B. A., Griffin-Jones, C., & Canfield, A. E. (2004) Chondrogenic and adipogenic potential of microvascular pericytes Circulation 110: 2226-2232. doi:10.1161/01.CIR.0000144457.55518.E5

Gilsanz, V., Kremer, A., Mo, A. O., Wren, T. A., & Kremer, R. (2010) Vitamin D status and its relation to muscle mass and muscle fat in young women J Clin Endocrinol Metab 95: 1595-1601. doi:10.1210/jc.2009-2309

Groenen, M. A. M., Archibald, A. L., Uenishi, H., Tuggle, C. K., Takeuchi, Y., Rothschild, M. F., . . . Schook, L. B. (2012) Analyses of pig genomes provide insight into porcine demography and evolution Nature 491: 393-398. doi:10.1038/nature11622

Handschin, C., & Spiegelman, B. M. (2011) PGC-1 coactivators and the regulation of skeletal muscle fiber-type determination Cell Metab 13: 351. doi:10.1016/j.cmet.2011.03.008

Hausman, G. J., Basu, U., Du, M., Fernyhough-Culver, M., & Dodson, M. V. (2014) Intermuscular and intramuscular adipose tissues: Bad vs. good adipose tissues Adipocyte 3: 242-255. doi:10.4161/adip.28546

He, Y. F., Xu, K., Li, Y. F., Chang, H., Liao, X., Yu, H., . . . Shi, L. (2022) Metabolomic Changes Upon Conjugated Linoleic Acid Supplementation and Predictions of Body Composition Responsiveness J Clin Endocrinol Metab 107: 2606-2615. doi:10.1210/clinem/dgac367

Hommelberg, P. P. H., Plat, J., Remels, A. H. V., van Essen, A. L. M., Kelders, M. C. J. M., Mensink, R. P., . . . Langen, R. C. J. (2010) Trans-10, cis-12 conjugated linoleic acid inhibits skeletal muscle differentiation and GLUT4 expression independently from NF-kappa B activation Mol Nutr Food Res 54: 1763-1772. doi:10.1002/mnfr.201000103

Jang, Y. J., Koo, H. J., Sohn, E. H., Kang, S. C., Rhee, D. K., & Pyo, S. (2015) Theobromine inhibits differentiation of 3T3-L1 cells during the early stage of adipogenesis via AMPK and MAPK signaling pathways Food Funct 6: 2365-2374. doi:10.1039/c5fo00397k

Jiang, Z., Marriott, K., & Maly, M. R. (2019) Impact of Inter-and Intramuscular Fat on Muscle Architecture and Capacity Crit Rev Biomed Eng 47: 515-533. doi:10.1615/CritRevBiomedEng.2020031124

Jin, L., Tang, Q., Hu, S., Chen, Z., Zhou, X., Zeng, B., . . . Li, M. (2021) A pig BodyMap transcriptome reveals diverse tissue physiologies and evolutionary dynamics of transcription Nat Commun 12: 3715. doi:10.1038/s41467-021-23560-8

Joe, A. W., Yi, L., Natarajan, A., Le Grand, F., So, L., Wang, J., . . . Rossi, F. M. (2010) Muscle injury activates resident fibro/adipogenic progenitors that facilitate myogenesis Nat Cell Biol 12: 153-163. doi:10.1038/ncb2015

Lang, I., Schweizer, A., Hiden, U., Ghaffari-Tabrizi, N., Hagendorfer, G., Bilban, M., . . . Desoye, G. (2008) Human fetal placental endothelial cells have a mature arterial and a juvenile venous phenotype with adipogenic and osteogenic differentiation potential Differentiation 76: 1031-1043. doi:10.1111/j.1432-0436.2008.00302.x

Lunney, J. K., Van Goor, A., Walker, K. E., Hailstock, T., Franklin, J., & Dai, C. H. (2021) Importance of the pig as a human biomedical model Sci Transl Med 13: eabd5758. doi:10.1126/scitranslmed.abd5758

M, K. K., S, I., & M, B. K. (2013) Mice do not accumulate muscle lipid in response to dietary conjugated linoleic acid. J Anim Sci 91: 4705–4712.

Men, X. M., Deng, B., Xu, Z. W., Tao, X., & Qi, K. K. (2013) Age-related changes and nutritional regulation of myosin heavy-chain composition in longissimus dorsi of commercial pigs Animal 7: 1486-1492. doi:10.1017/S1751731113000992

Mitchell, K. J., Pannerec, A., Cadot, B., Parlakian, A., Besson, V., Gomes, E. R., . . . Sassoon, D. A. (2010) Identification and characterization of a non-satellite cell muscle resident progenitor during postnatal development Nat Cell Biol 12: 257-U256. doi:10.1038/ncb2025

Parra, P., Serra, F., & Palou, A. (2012) Transcriptional analysis reveals a high impact of conjugated linoleic acid on stearoyl-Coenzyme A desaturase 1 mRNA expression in mice gastrocnemius muscle Genes Nutr 7: 537-548. doi:10.1007/s12263-011-0279-x

Petrany, M. J., Swoboda, C. O., Sun, C., Chetal, K., Chen, X., Weirauch, M. T., . . . Millay, D. P. (2020) Single-nucleus RNA-seq identifies transcriptional heterogeneity in multinucleated skeletal myofibers Nat Commun 11: 6374. doi:10.1038/s41467-020-20063-w

Sahlin, K. R., & Trewern, J. (2022) A systematic review of the definitions and interpretations in scientific literature of ’less but better’ meat in high-income settings Nature Food 3: 454-460. doi:10.1038/s43016-022-00536-5

Schiaffino, S., & Reggiani, C. (2011) Fiber types in mammalian skeletal muscles Physiol Rev 91: 1447-1531. doi:10.1152/physrev.00031.2010

Sciorati, C., Clementi, E., Manfredi, A. A., & Rovere-Querini, P. (2015) Fat deposition and accumulation in the damaged and inflamed skeletal muscle: cellular and molecular players Cell Mol Life Sci 72: 2135-2156. doi:10.1007/s00018-015-1857-7

Shan, T., Xiong, Y., Zhang, P., Li, Z., Jiang, Q., Bi, P., . . . Kuang, S. (2016) Lkb1 controls brown adipose tissue growth and thermogenesis by regulating the intracellular localization of CRTC3 Nat Commun 7: 12205. doi:10.1038/ncomms12205

Tabula Muris, C., Overall, C., Logistical, C., Organ, C., processing, Library, P., . . . Principal, I. (2018) Single-cell transcriptomics of 20 mouse organs creates a Tabula Muris Nature 562: 367-372. doi:10.1038/s41586-018-0590-4

Tamaki, T., Akatsuka, A., Ando, K., Nakamura, Y., Matsuzawa, H., Hotta, T., . . . Edgerton, V. R. (2002) Identification of myogenic-endothelial progenitor cells in the interstitial spaces of skeletal muscle J Cell Biol 157: 571-577. doi:10.1083/jcb.200112106

Terasawa, N., Okamoto, K., Nakada, K., & Masuda, K. (2017) Effect of Conjugated Linoleic Acid Intake on Endurance Exercise Performance and Anti-fatigue in Student Athletes J Oleo Sci 66: 723-733. doi:10.5650/jos.ess17053

Uezumi, A., Fukada, S., Yamamoto, N., Takeda, S., & Tsuchida, K. (2010) Mesenchymal progenitors distinct from satellite cells contribute to ectopic fat cell formation in skeletal muscle Nat Cell Biol 12: 143-152. doi:10.1038/ncb2014

van Vliet, S., Fappi, A., Reeds, D. N., & Mittendorfer, B. (2020) No independent or combined effects of vitamin D and conjugated linoleic acids on muscle protein synthesis in older adults: a randomized, double-blind, placebo-controlled clinical trial Am J Clin Nutr 112: 1382-1389. doi:10.1093/ajcn/nqaa240

Wang, L., Huang, Y., Wang, Y., & Shan, T. (2021) Effects of Polyunsaturated Fatty Acids Supplementation on the Meat Quality of Pigs: A Meta-Analysis Front Nutr 8: 746765. doi:10.3389/fnut.2021.746765

Wang, L., Nong, Q., Zhou, Y., Sun, Y., Chen, W., Xie, J., . . . Shan, T. (2022) Changes in Serum Fatty Acid Composition and Metabolome-Microbiome Responses of Heigai Pigs Induced by Dietary N-6/n-3 Polyunsaturated Fatty Acid Ratio Front Microbiol 13: 917558. doi:10.3389/fmicb.2022.917558

Wang, L., Valencak, T. G., & Shan, T. (2024) Fat infiltration in skeletal muscle: Influential triggers and regulatory mechanism iScience 27: 109221. doi:10.1016/j.isci.2024.109221

Wang, L., Zhang, S., Huang, Y., You, W., Zhou, Y., Chen, W., . . . Shan, T. (2022) CLA improves the lipo-nutritional quality of pork and regulates the gut microbiota in Heigai pigs Food Funct 12093-12104. doi:10.1039/d2fo02549c

Wang, L., Zhang, S., Huang, Y., Zhou, Y., & Shan, T. (2023) Conjugated linoleic acids inhibit lipid deposition in subcutaneous adipose tissue and alter lipid profiles in serum of pigs J Anim Sci 101: skad294. doi:10.1093/jas/skad294

Wang, L., Zhao, X., Liu, S., You, W., Huang, Y., Zhou, Y., . . . Shan, T. (2023) Single-nucleus and bulk RNA sequencing reveal cellular and transcriptional mechanisms underlying lipid dynamics in high marbled pork NPJ Sci Food 7: 23. doi:10.1038/s41538-023-00203-4

Wei, H., Zhou, Y., Jiang, S., Huang, F., Peng, J., & Jiang, S. (2016) Transcriptional response of porcine skeletal muscle to feeding a linseed-enriched diet to growing pigs J Anim Sci Biotechnol 7: 6. doi:10.1186/s40104-016-0064-1

Wosczyna, M. N., Perez Carbajal, E. E., Wagner, M. W., Paredes, S., Konishi, C. T., Liu, L., . . . Rando, T. A. (2021) Targeting microRNA-mediated gene repression limits adipogenic conversion of skeletal muscle mesenchymal stromal cells Cell Stem Cell 28: 1323-1334 e1328. doi:10.1016/j.stem.2021.04.008

Xu, D. D., Wan, B. Y., Qiu, K., Wang, Y. B., Zhang, X., Jiao, N., . . . Yin, J. D. (2023) Single-Cell RNA-Sequencing Provides Insight into Skeletal Muscle Evolution during the Selection of Muscle Characteristics Advanced Science. doi:10.1002/advs.202305080

Xu, Z., You, W., Chen, W., Zhou, Y., Nong, Q., Valencak, T. G., . . . Shan, T. (2020) SingleDcell RNA sequencing and lipidomics reveal cell and lipid dynamics of fat infiltration in skeletal muscle Journal of Cachexia, Sarcopenia and Muscle 12: 109–129. doi:10.1002/jcsm.12643

Yamakawa, D., Tsuboi, J., Kasahara, K., Matsuda, C., Nishimura, Y., Kodama, T., . . . Inagaki, M. (2023) Cilia-Mediated Insulin/Akt and ST2/JNK Signaling Pathways Regulate the Recovery of Muscle Injury Adv Sci 10: e2202632. doi:10.1002/advs.202202632

Yeganeh, A., Taylor, C. G., Poole, J., Tworek, L., & Zahradka, P. (2016) Trans10, cis12 conjugated linoleic acid inhibits 3T3-L1 adipocyte adipogenesis by elevating beta-catenin levels Biochimica Et Biophysica Acta-Molecular and Cell Biology of Lipids 1861: 363–370. doi:10.1016/j.bbalip.2016.01.004

Yi, J. M., Zhu, J. J., Wu, J., Thompson, C. B., & Jiang, X. J. (2020) Oncogenic activation of PI3K-AKT-mTOR signaling suppresses ferroptosis via SREBP-mediated lipogenesis Proc Natl Acad Sci U S A 117: 31189-31197. doi:10.1073/pnas.2017152117

Yi, W., Huang, Q., Wang, Y., & Shan, T. (2023) Lipo-nutritional quality of pork: The lipid composition, regulation, and molecular mechanisms of fatty acid deposition Anim Nutr 13: 373-385. doi:10.1016/j.aninu.2023.03.001

Yin, H., Pasut, A., Soleimani, V. D., Bentzinger, C. F., Antoun, G., Thorn, S., . . . Rudnicki, M. A. (2013) MicroRNA-133 controls brown adipose determination in skeletal muscle satellite cells by targeting Prdm16 Cell Metab 17: 210-224. doi:10.1016/j.cmet.2013.01.004

You, W. J., Liu, S. Q., Ji, J. F., Ling, D. F., Tu, Y., Zhou, Y. B., . . . Shan, T. Z. (2023) Growth arrest and DNA damage-inducible alpha regulates muscle repair and fat infiltration through ATP synthase F1 subunit alpha Journal of Cachexia Sarcopenia and Muscle 14: 326-341. doi:10.1002/jcsm.13134

Zhang, H., Dong, X., Wang, Z., Zhou, A., Peng, Q., Zou, H., . . . Wang, L. (2016) Dietary conjugated linoleic acids increase intramuscular fat deposition and decrease subcutaneous fat deposition in Yellow Breed x Simmental cattle Anim Sci J 87: 517-524. doi:10.1111/asj.12447

Zhao, L., Son, J. S., Wang, B., Tian, Q., Chen, Y., Liu, X., . . . Du, M. (2020) Retinoic acid signalling in fibro/adipogenic progenitors robustly enhances muscle regeneration EBioMedicine 60: 103020. doi:10.1016/j.ebiom.2020.103020

Zhao, S. H., Cheng, L., Shi, Y., Li, J., Yun, Q. H., & Yang, H. (2021) MIEF2 reprograms lipid metabolism to drive progression of ovarian cancer through ROS/AKT/mTOR signaling pathway Cell Death Dis 12: 18. doi:10.1038/s41419-020-03336-6

