## supplementary materials for "Single-nucleus transcriptomics reveal the cytological mechanism of conjugated linoleic acids in regulating intramuscular fat deposition"

**Table S1.** The primer sequence of qPCR.

| Primer name | Sequence (5'-3') |
| --- | --- |
| 18s-F | CCCACGGAATCGAGAAAGAG |
| 18s-R | TTGACGGAAGGGCACCA |
| ACACA-F | AGCAAGGTCGAGACCGAAAG |
| ACACA-R | TAAGACCACCGGCGGATAGA |
| ADIPOQ-F | CTCCTTCCACGTCACGGTCT |
| ADIPOQ-R | CCAGATAGAGGAGCACAGAGCC |
| ANO4-F | GGTCTGAATCGTCTGCTTACTAATGG |
| ANO4-R | TCCCTTGTGAAGTCCTTTCCTAGAG |
| ARBB1-F | GAACTCCGTGCGTCTGGTCATC |
| ARBB1-R | AGGAACTGCCTGGTGGTCTCG |
| ATGL-F | GCACCTTCATTCCCGTGTAC |
| ATGL-R | TTGTCTGAGATGCCACCGTC |
| COX1-F | AACTGACTCGTACCGCTAATAATCG |
| COX1-R | GGATGCCAGAAGTAATAGGAAGGATG |
| DGAT2-F | AGGACATTGACCTCTACCATGC |
| DGAT2-R | CAGTTCACCTCCAGGACCTC |
| ELOVL6-F | AGAACACGTAGCGACTCCGAAGAT |
| ELOVL6-R | GACATGCCGACCGCCAAAGATAA |
| FABP4-F | TGGAAACTTGTCTCCAGTG |
| FABP4-R | GGTACTTTCTGATCTAATGGTG |
| FABP5-F | ACTGTCTGCGACTTTACCAATGG |
| FABP5-R | TTCTTGTGATTGTGCTCTCCTTCC |
| FASN-F | GCAGGCGCGTGATGGGAATGGTG |
| FASN-R | GCCCGAGCCCGAGTGGATGAGCA |
| HSL-F | CCCCCGTGCGCTGGAGGAGT |
| HSL-R | GGGAGGGGGAGGCGGCAGAC |
| MYBPC1-F | CTATTCTCAGCCCATTCTCGTG |
| MYBPC1-R | TCTGGTCTTGGTTTTCCCTG |
| NEB-F | AGGAAGCAATAGGACAAGGAAC |
| NEB-R | CAATCTCTGGAGTCACAGTGG |
| PDE4D-F | GGAAGATGGCGAGTCAGATACG |
| PDE4D-R | TGGCTCTCCTCCTCCTCTCC |
| PDE7B-F | CCTACATCGTGGAGCCACTCTTC |
| PDE7B-R | TGCTACCGCTGCTGCTTCTG |
| PPARγ-F | GGCGAGGGCGATCTTGACAG |
| PPARγ-R | GATGCGAATGGCCACCTCTTT |
| SCD-F | CAAACACCCAGCCGTCAAAG |
| SCD-R | CGAAGAAAGGTGGCGACGAA |
| SIAH1-F | TTAATCTTCCTGGTGCTGTTGACTG |
| SIAH1-R | GCTTGCTTGCGTGTTCCTATCAG |
| SREBP1-F | GTGCTGGCGGAGGTCTATGT |
| SREBP1-R | AGGAAGAAGCGGGTCAGAAAG |
| THBS1-F | CTGGACTTGCTGTAGGTTATGATGAG |
| THBS1-R | CATAGAAACGGCTGCTGGACTG |
| TIMP3-F | CCTTTGGCACACTGGTCTACAC |
| TIMP3-R | GGTACTGGTACTTGTTGACTTCTAGC |

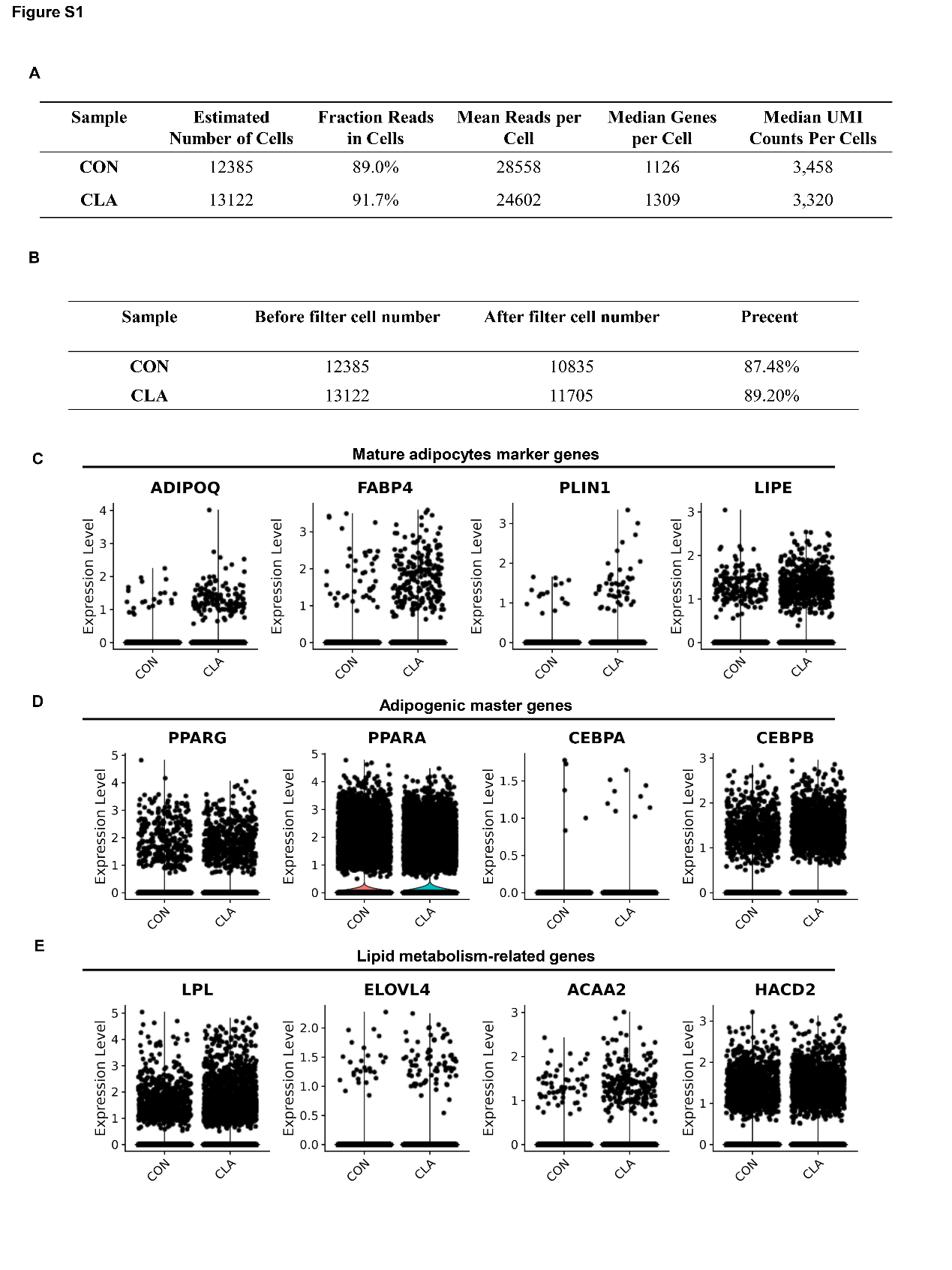

**Supplementary Figure 1. CLA upregulated the expression of adipogenic related genes in muscles.** **(A)** The results obtained from Cell Ranger analyses. **(B)** Cell number of snRNA-seq datasets before or after filter from each sample. **(C-E)** Violin plot displaying the expression of adipogenic master genes **(C)**, mature adipocyte marker genes **(D)**, and lipid metabolism-related genes **(E)** in different groups.

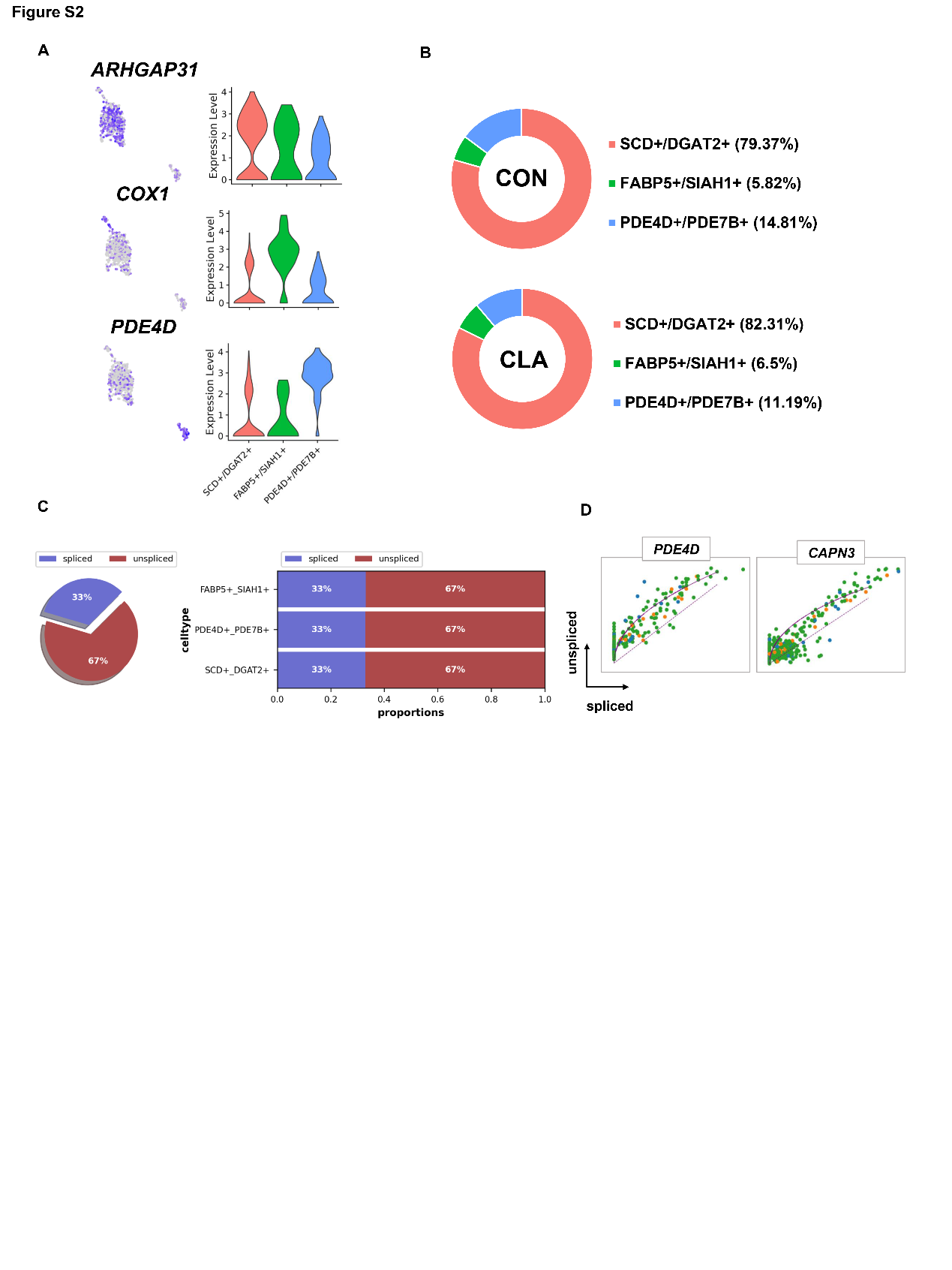

**Supplementary Figure 2.** **Clustering analysis of adipocytes nuclei. (A)** UMAP plot and violin plot showing the expression of three subcluster marker genes in adipocytes nuclei. **(B)** Cell proportion in each subcluster in different groups. Each cluster is colour-coded. **(C)** Pie chart showing the proportion of adipocytes that are spliced versus un-spliced. **(D)** Transcriptional dynamics of marker genes on the UMAPs based on RNA velocity analysis.

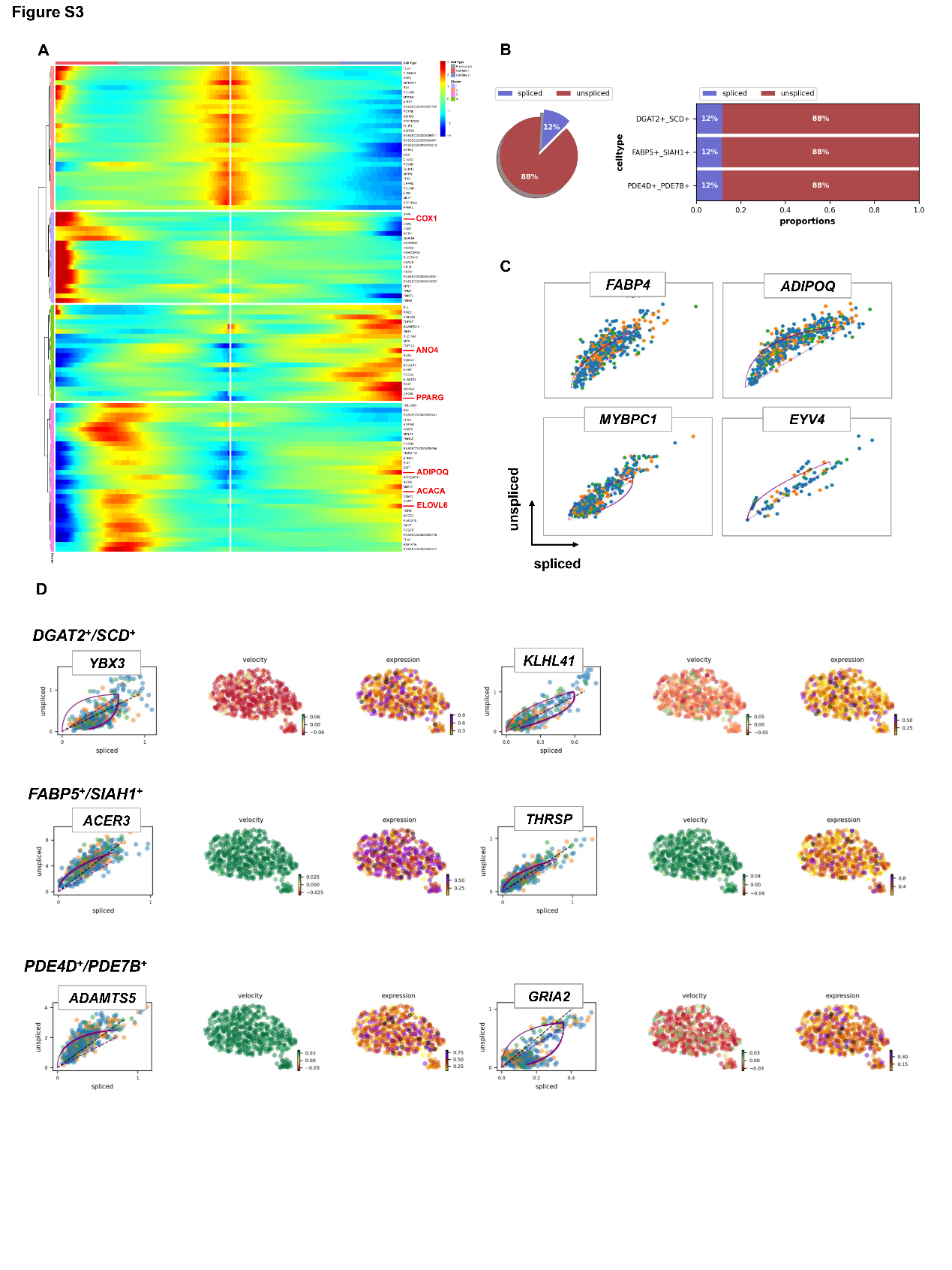

**Supplementary Figure 3.** **Pseudotime trajectory analysis of adipocytes nuclei by RNA velocity in HLW pigs. (A)** Pseudotemporal heatmap showing gene expression dynamics for significant marker genes. Genes (rows) were clustered into three modules, and cells (columns) were ordered according to pseudotime in different groups. **(B)** Pie chart showing the proportion of FAPs subtypes that are spliced versus un-spliced. **(C)** Transcriptional dynamics of marker genes on the UMAPs based on RNA velocity analysis. **(D)** Transcriptional dynamics of top 2 marker genes on the UMAPs based on RNA velocity analysis.

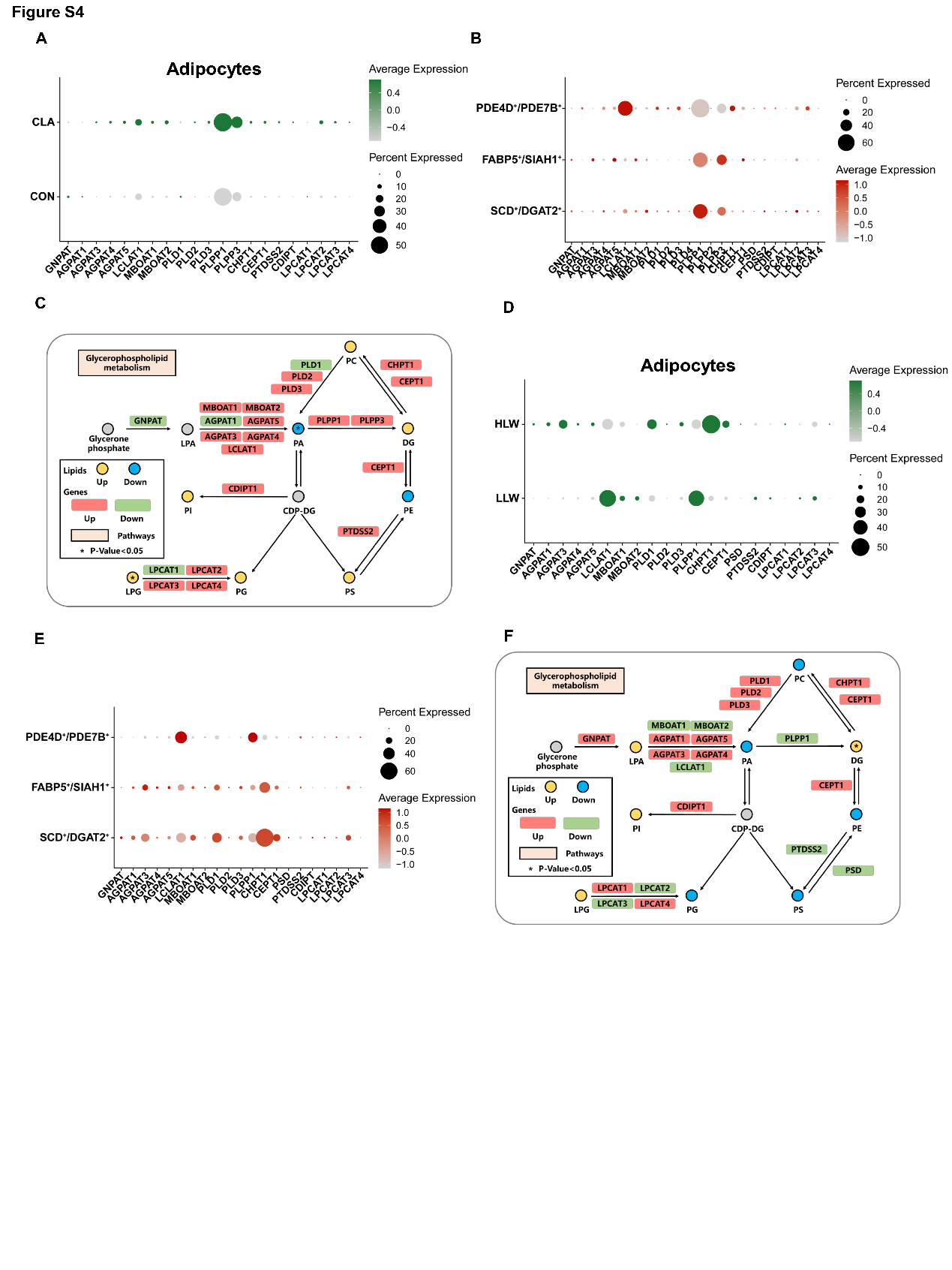

**Supplementary Figure 4.** **Comparison of gene programs involved in glycerophospholipid metabolism. (A)** Dot plot showing the relative expression of glycerophospholipid metabolism-related genes in CON and CLA group of adipocytes. **(B)** Dot plot showing the relative expression of glycerophospholipid metabolism-related genes in three subclusters of adipocytes nuclei in Heigai pigs. **(C)** Selected glycerophospholipid metabolic reactions from KEGG, with indications of quantified lipid classes and acyl chains (circles) and genes (rectangles) significantly regulated compared with CON group. **(D)** Dot plot showing the relative expression of glycerophospholipid metabolism-related genes in HLW and LLW group of adipocytes nuclei. **(E)** Dot plot showing the relative expression of glycerophospholipid metabolism-related genes in three subclusters of adipocytes nuclei in Laiwu pigs. **(F)** Selected glycerophospholipid metabolic reactions from KEGG, with indications of quantified lipid classes and acyl chains (circles) and genes (rectangles) significantly regulated compared with LLW group. Statistical analysis was performed using two-tailed Student’s t-test.

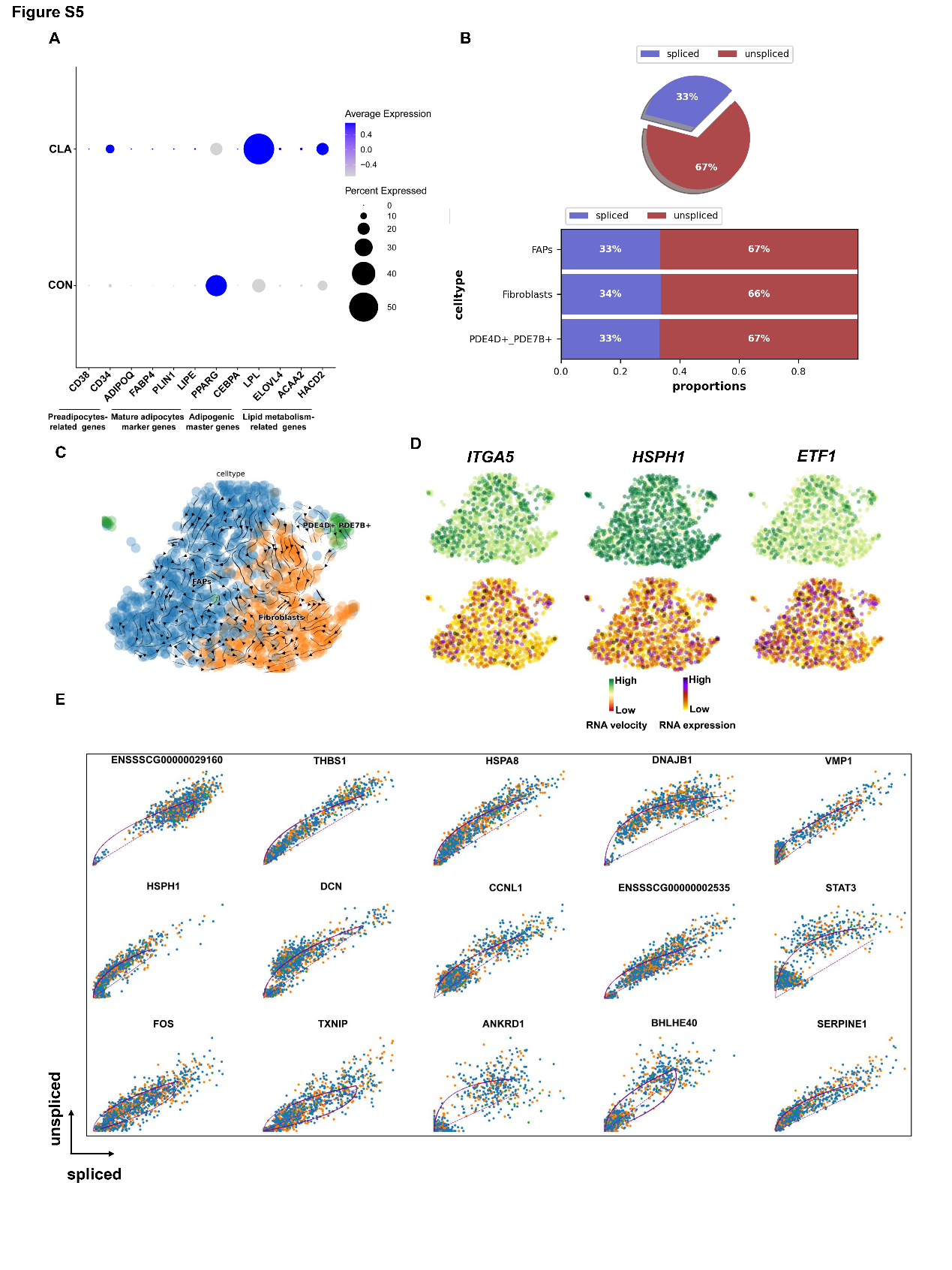

**Supplementary Figure 5.** **Pseudotime trajectory analysis of FAPs nuclei by RNA velocity. (A)** Dot plot showing the expression of preadipocytes-related genes (*CD38*, and *CD34*), mature adipocytes marker genes (*ADIPOQ*, *FABP4*, *PLIN1* and *LIPE*), adipogenic master (*PPARG* and *CEBPA*), and lipid metabolism-related genes (*LPL*, *ELOVL4*, *ACAA2* and *HACD2*) after CLA treatment in FAPs. **(B)** Pie chart showing the proportion of adipocytes that are spliced versus un-spliced. Bar plot showing the proportion of FAPs subtypes that are spliced versus un-spliced in different species. **(C)** Unsupervised pseudotime trajectory of the three subtypes of FAPs by RNA velocity analysis. Trajectory is colored by cell subtypes. The arrow indicates the direction of cell pseudo-temporal differentiation. **(D)** Distribution of marker genes of the three subtypes on the UMAPs based on RNA velocity analysis. **(E)** Transcriptional dynamics of top 15 marker genes on the UMAPs based on RNA velocity analysis.

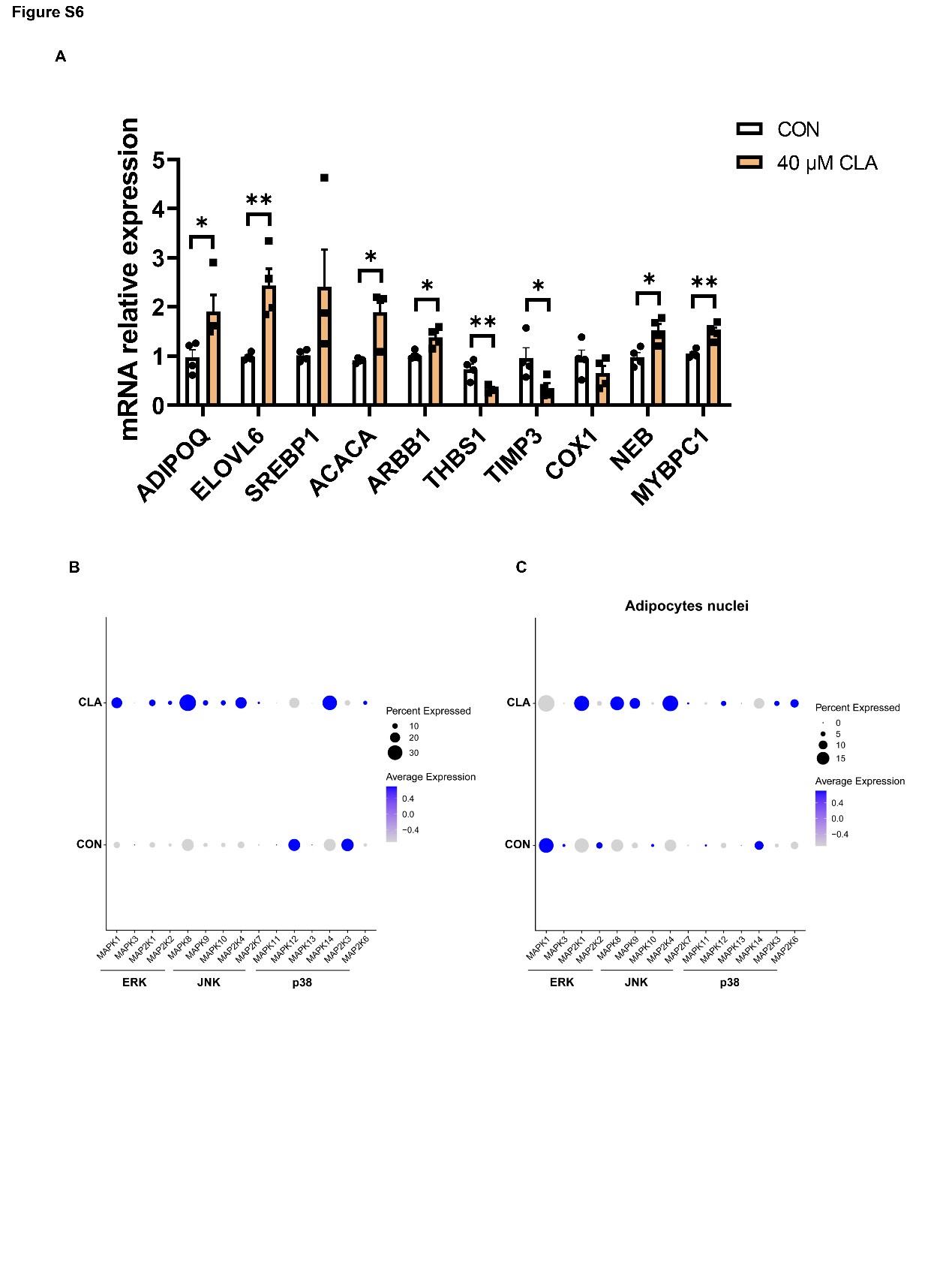

**Supplementary Figure 6. Changes of MAPK signalling pathway in muscle nuclei. (A)** The mRNA expression of marker genes at key points of FAPs differentiation (n=4). **(B-C)** Dot plot showing the expression of MAPK signalling pathway related genes after CLA treatment in muscle **(B)** and adipocytes **(C)**. Error bars represent SEM. **P* < 0.05, ** *P* < 0.01, two-tailed Student’s t-test.
